# Structural basis for the Pr-Pfr long-range signaling mechanism of a full-length bacterial phytochrome at the atomic level

**DOI:** 10.1101/2021.02.16.430237

**Authors:** Lisandro H. Otero, Sabrina Foscaldi, Giuliano T. Antelo, Serena Sirigu, Sebastián Klinke, Lucas A. Defelipe, Maximiliano Sánchez-Lamas, Giovanni Battocchio, Leonard M. G. Chavas, Fernando A. Goldbaum, Maria-Andrea Mroginski, Jimena Rinaldi, Hernán R. Bonomi

## Abstract

Light sensing allows organisms to adapt to constantly changing environmental factors. Phytochromes constitute a widespread biological photoreceptor family that typically interconvert between two photostates called Pr (red light-absorbing) and Pfr (far-red light-absorbing). Despite the vast structural information reported on phytochromes, the lack of full-length structures at the (near-)atomic level in both pure Pr and Pfr states leaves gaps in the structural mechanisms involved in the signal transmission pathways during the photoconversion. Here we present three crystallographic structures from the plant pathogen *Xanthomonas campestris* virulence regulator bacteriophytochrome, including two full-length proteins, in the Pr and Pfr states. The structural findings, combined with mutational, biochemical and computational studies, allow us to describe the signaling mechanism of a full-length bacterial phytochrome at the atomic level, from the isomerization of the chromophore and the β-sheet/α-helix tongue transition to the remodeling of the quaternary assembly of the protein.

**TEASER:** Crystal structures of the full-length bacteriophytochrome *Xcc*BphP in both Pr and Pfr states unveil photoswitching mechanism

## INTRODUCTION

Light sensing mechanisms allow prokaryotes and eukaryotes to gather information about their ever-changing environments. Phytochromes are red and far-red light-sensing proteins that constitute a widespread biological photoreceptor family found in plants, algae, fungi, and prokaryotes (*1*, *2*). Interestingly, bacterial phytochromes (bacteriophytochromes, BphPs) have gained attention since their discovery (*3*, *4*) with growing numbers of studies addressing their roles in infective and pathogenic organisms.

The prototypical BphP architecture consists of an N-terminal photosensory module (PSM), which autocatalytically binds biliverdin IXα (BV) chromophore to a conserved cysteine residue, and a C-terminal variable output module (OM), responsible for transducing the PSM light-driven conformational changes into a specific physiological signal (*5*–*9*).

The PSM is typically composed of three modular domains linearly structured sharing topological elements: a Per-Arnt-Sim domain (PAS), followed by cyclic GMP-Adenylyl cyclase-FhlA (GAF) and phytochrome-specific (PHY) domains. The OM is often composed of histidine kinase modules (HK) that are assumed to trigger a two-component signaling pathway *via* a phosphotransfer mechanism to regulate downstream processes such as gene expression. However, other frequent OM architectures include (i) HK–response regulator pairs, (ii) c-di-GMPcyclase/phosphodiesterases, and (iii) PAS domains, among others, evidencing the wide range of signaling outputs in the phytochrome family (*8*). Remarkably, the PSM-containing phytochromes presenting at least one PAS domain within the OM represent ~26% of the total sequences from the Pfam database (https://pfam.xfam.org/).

BphPs usually show a dimeric quaternary structure, where monomers are docked to each other in a parallel or antiparallel arrangement through long helical bundles across the different domains called GAF-PHY and PHY-OM helical linkers, which form the “helical spine” (*6*, *7*). Other outstanding structural features from these photoreceptors are (i) the presence of a figure-of-eight knot that crosses over residues between the PAS and GAF domains, strengthening their association (*10*), and (ii) a highly conserved hairpin protrusion from the PHY domain, termed as the “tongue”, which encloses the chromophore binding pocket by interacting electrostatically with the GAF domain (*6*–*9*).

The basic principle of BphP photochemistry is their reversible photoconversion, typically, between two photostates that exhibit different absorption spectra named Pr (red light-absorbing) and Pfr (far-red light-absorbing) (*11*, *12*). In the dark, canonical type BphPs exhibit a Pr thermal ground state, while bathy type BphPs show a Pfr thermal ground state (*13*). Although the full picture on how the light-induced changes are transduced from the PSM to the OM is still incomplete, there are a series of well-established reversible structural features during the Pr/Pfr photoconversion (*14*–*24*).

Upon red (far-red) light absorption in the Pr (Pfr) state, the initial step of the intramolecular signaling mechanism, namely the BV chromophore photoisomerization, is triggered. This initial step results in the Lumi-R (Lumi-F) intermediate which involves a *Z/E* (*E/Z*) conversion of the C15 = C16 double bond between the pyrrole rings C and D along with a series of transient proto translocation (Meta-R or Meta-F intermediates) events at the rings B and C pyrrole nitrogen atoms, as well as protonation dynamic events in the biliverdin binding pocket. As a result, a ~180° rotation of the ring D is produced, defining a *ZZZssa* or *ZZEssa* BV configuration in the Pr or Pfr states, respectively. The time-scale of the *ZZZssa*-to-*ZZEssa* intermediate formation ranges from ultrafast transitions ranging in the picosecond range (Lumi intermediates) towards microseconds (Meta species). The distinct intermediate states have been elucidated in view of their structural and spectroscopic features (*25*–*28*).

As the chromophore interacts with residues located at different regions of the PSM, the local physicochemical changes upon photoisomerization perturb the structure of the PSM. Consequently, the “tongue” reversibly interconverts into two structural conformations, a two-stranded antiparallel β-sheet in the Pr state, and an α-helix in the Pfr state (*15*, *18*–*24*). Although it is not currently fully understood how the tongue structural changes develop, it is well agreed that the β-sheet/α-helix tongue transition generates a push/pull movement between the GAF and PHY domains, reorienting the GAF-PHY and PHY-OM helical linkers from the helical spine, and thus, modifying the dimer interface (*22*, *24*, *29*). This large-scale structural motion is proposed to be critical for modulating the OM signaling activity. Nevertheless, the structural determinants of this mechanism of photoreception are still poorly understood as most of the BphP structures reported to date lack their corresponding OMs and, additionally, no full-length phytochrome structures have been solved at the (near-)atomic level in both pure (non-mixed) Pr and Pfr photostates (*9*).

Our group studies the BphP from the plant pathogen *Xanthomonas campestris* pv. *campestris* (*Xcc*BphP), which functions as virulence regulator (*30*). *Xcc*BphP has been classified as a bathy-like phytochrome as it normally thermally decays to a mixed Pfr:Pr equilibrium in the dark (~6:1) upon irradiation (*31*). Moreover, we have solved the crystal structure of the full-length version bearing its PSM (formed by the domain triad PAS2-GAF-PHY) as well as its complete OM (a PAS9 domain) in the Pr state at 3.25 Å resolution (*31*, *32*). However, the structure showed weak electron density at the region corresponding to ring D of the BV molecule, so neither a *ZZZssa* nor a *ZZEssa* chromophore configuration could be defined based solely on the X-ray crystallographic data (*31*). Due to the lack of a *Xcc*BphP Pfr structure, we used the *Rhodopseudomonas palustris Rp*BphP1 Pfr structure (sharing a similar domain architecture) (*33*) to propose a rough model for signal propagation from the PSM to the OM during the *Xcc*BphP light-driven conversion mechanism. However, the use of two different BphPs to build a precise mechanistic intramolecular transducing model at the atomic scale is far from optimal.

In this study, we present two novel crystal structures of *Xcc*BphP variants in the Pfr state, including a full-length version. In addition, we have obtained a new wild-type full-length Pr structure with an enhanced resolution with the complete chromophore molecule clearly presenting a *ZZZssa* configuration. These structures enabled us to identify unprecedented structural rearrangements between both photostates in a complete BphP. Spectroscopic and mutational analyses along with computational modeling and biophysical experiments have allowed us to probe the mechanism at play. Our results enabled us to build a more complete model on the light-driven conformational changes transmitted from the PSM to the OM in the phytochrome photocycle.

## RESULTS

### Rationale and strategy

To get insights into the signaling mechanisms at the atomic level during the reversible Pr-Pfr photoswitching in full-length BphPs, we aimed at solving the crystal structures of *Xcc*BphP variants in the Pr and Pfr photostates.

First, one of our aims was to obtain a better crystal structure of the Pr form previously solved to clearly define the BV ring D, and unambiguously allocate a *ZZZssa* chromophore configuration. Regarding the Pfr form, despite great efforts, we failed to obtain this photostate in the crystal form using the wild-type version, even in dark conditions, where the highest Pfr proportion (~85%) is reached at equilibrium (*31*). In fact, in these conditions there is still a considerable Pr population from which we hypothesize crystals of *Xcc*BphP grow (*32*). Hence, our strategy to obtain the Pfr crystal structure was to use Pfr-favoured variants, for which the dark conversion rate is increased and/or the equilibrium is shifted.

We have previously demonstrated that a deletion of the C-terminal PAS9 domain, resulting in a shorter protein corresponding to the 1-511 residue range here named ΔPAS9(1-511), generates an active bathy type PSM with a much more relatively stable Pfr state (*31*). This is evidenced by a faster Pr-to-Pfr dark conversion kinetics and a higher Pfr:Pr ratio at thermal equilibrium. Additionally, in order to obtain a full-length Pfr-favored variant, we have recently designed a UV-Vis spectroscopy-based screening from clarified crude extracts containing overexpressed phytochromes to rapidly and simultaneously characterize the photochemistry of a large number of *Xcc*BphP variants. From this screening, the substitution G454E in the tongue region of the PHY domain was identified as the most Pfr-favoured candidate (*34*). Thus, both ΔPAS9(1-511) and G454E variants were chosen for crystallization trials.

### Crystallization and resolution of the *Xcc*BphP Pr and Pfr structural variants

Before the crystallization assays, we performed a brief characterization of protein properties in solution. The *Xcc*BphP wild-type and ΔPAS9(1-511) variants have been previously proven to assemble in the Pr form in approximately 5 min (*31*). In contrast, G454E shows a slower dark assembly also in the Pr form (**Figure S1A**). ΔPAS9(1-511) and G454E photoconvert upon red and far-red irradiation as the wild-type version (**Figure S1B**) (*31*). The dark conversion kinetics of these variants, and others that will be presented during this work, is analyzed in **Figures S2-S4** and summarized in **Figure 1** and **Table 1**. During dark conversion, both ΔPAS9(1-511) and G454E are enriched in the Pfr form compared to the wild-type version, reaching proportions over 96% of Pfr at equilibrium and with rapid kinetics, presenting half-life values of ~15 min and 50 min for G454E and ΔPAS9(1-511), respectively (the wild-type estimated half-life was ~307 min) **(Table 1**).

**Figure 1.**
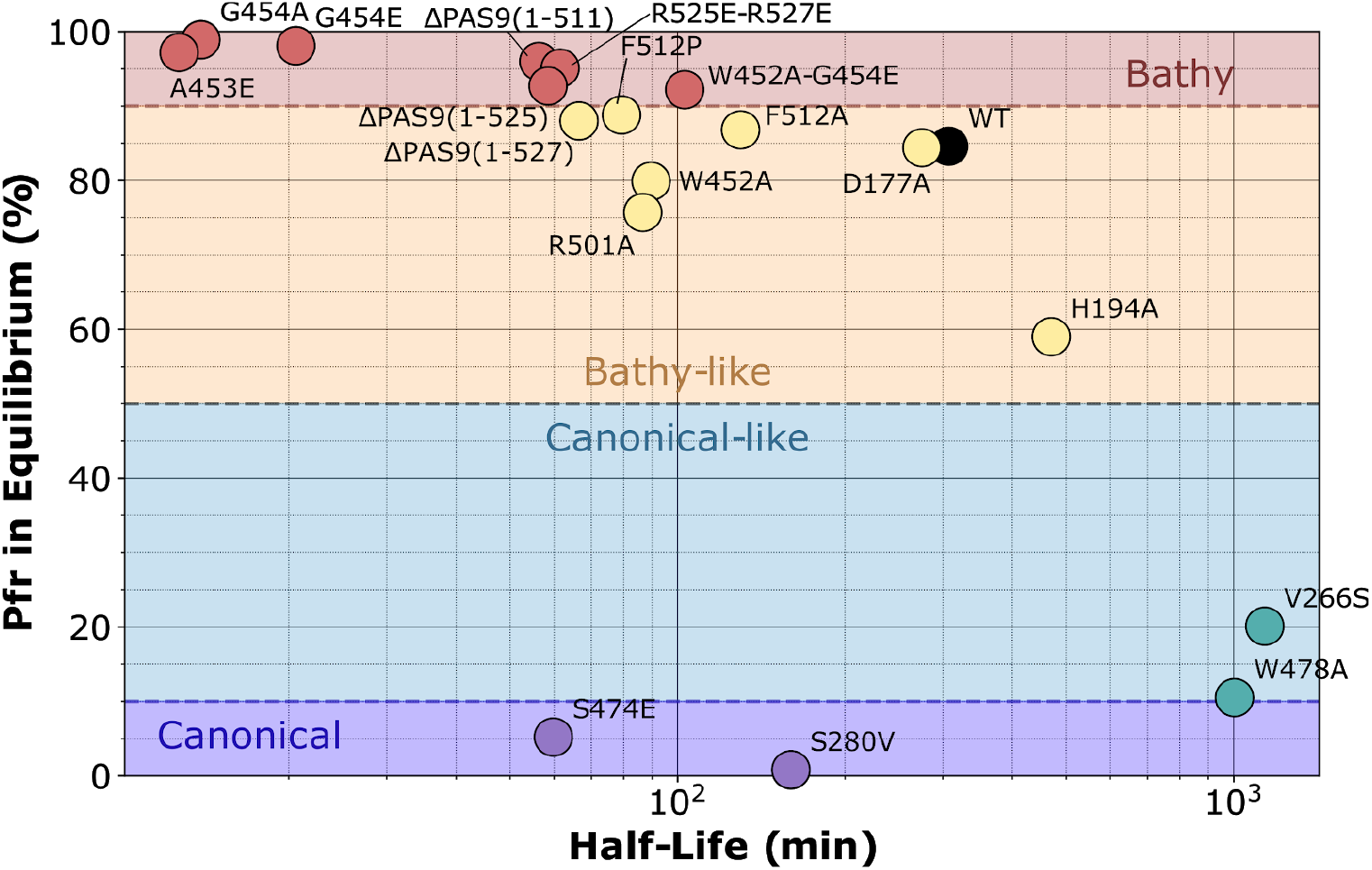
Kinetic parameters calculated for dark conversion of *Xcc*BphP variants. The photochemical behaviour of all variants studied in this work are summarized in this scatter plot, where the x-axis corresponds to the half-life values and the y-axis corresponds to the Pfr fractions (%) at equilibrium. These two parameters were estimated by fitting eq. 1 (monoexponential) and eq. 2 (biexponential) to the data from **Figure S2**, respectively (see also **Figure S3**). Each variant is represented by a different colored dot in the scatter plot. BphP categories were based on Pfr enrichment in equilibrium: bathy (>90%), bathy-like (50 to 90%), canonical-like (10 to 50%) and canonical (0 to 10%).

**Table 1.**
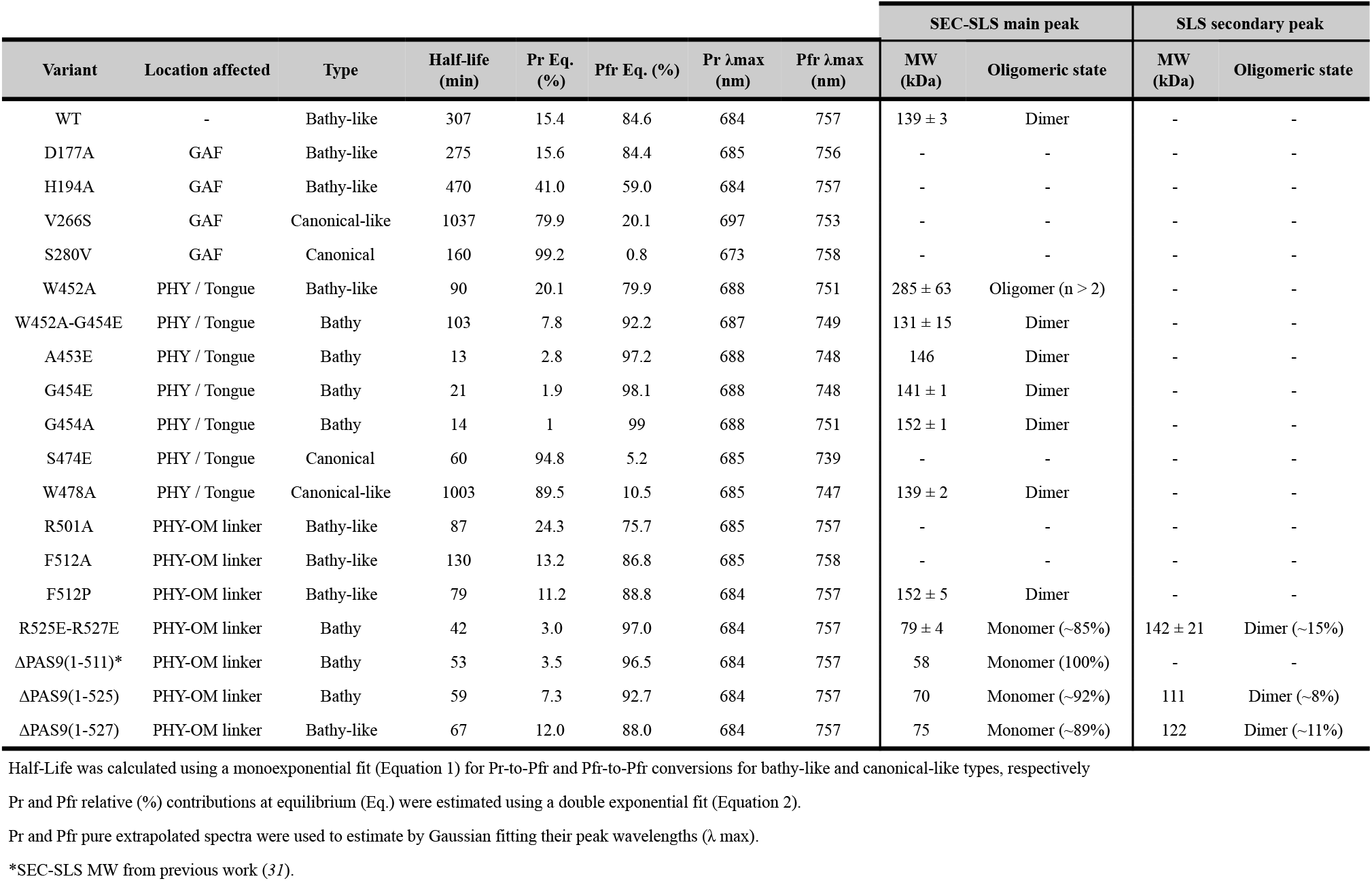
Kinetic and oligomerization parameters calculated for *Xcc*BPhP variants.

Wild-type *Xcc*BphP was crystallized following the reported protocol for the Pr state structure (*32*) in the same *P*4_3_2_1_2 space group and with equivalent unit cell parameters (**Table S2**). The G454E and ΔPAS9(1-511) variants crystallized in two different conditions (see Materials and Methods) in the *I*4_1_ space group.

The three crystallographic structures were solved by molecular replacement using the previously reported wild-type Pr structure (PDB entry 5AKP) as starting model and refined at 2.68 (G454E), 2.95 [ΔPAS9(1-511)] and 2.96 Å (wild-type protein) with favorable stereochemistry and good refinement statistics (**Table S2**). The final refined models show two chains (A and B) in the asymmetric unit in the wild-type, and only one in the G454E and ΔPAS9(1-511) variants (**Figure 2**). While the wild-type structure shows an overall fold characteristic of the Pr state, both the G454E and ΔPAS9(1-511) structures reveal a scaffold in total agreement with a Pfr state, as described in detail in the following sections.

**Figure 2.**
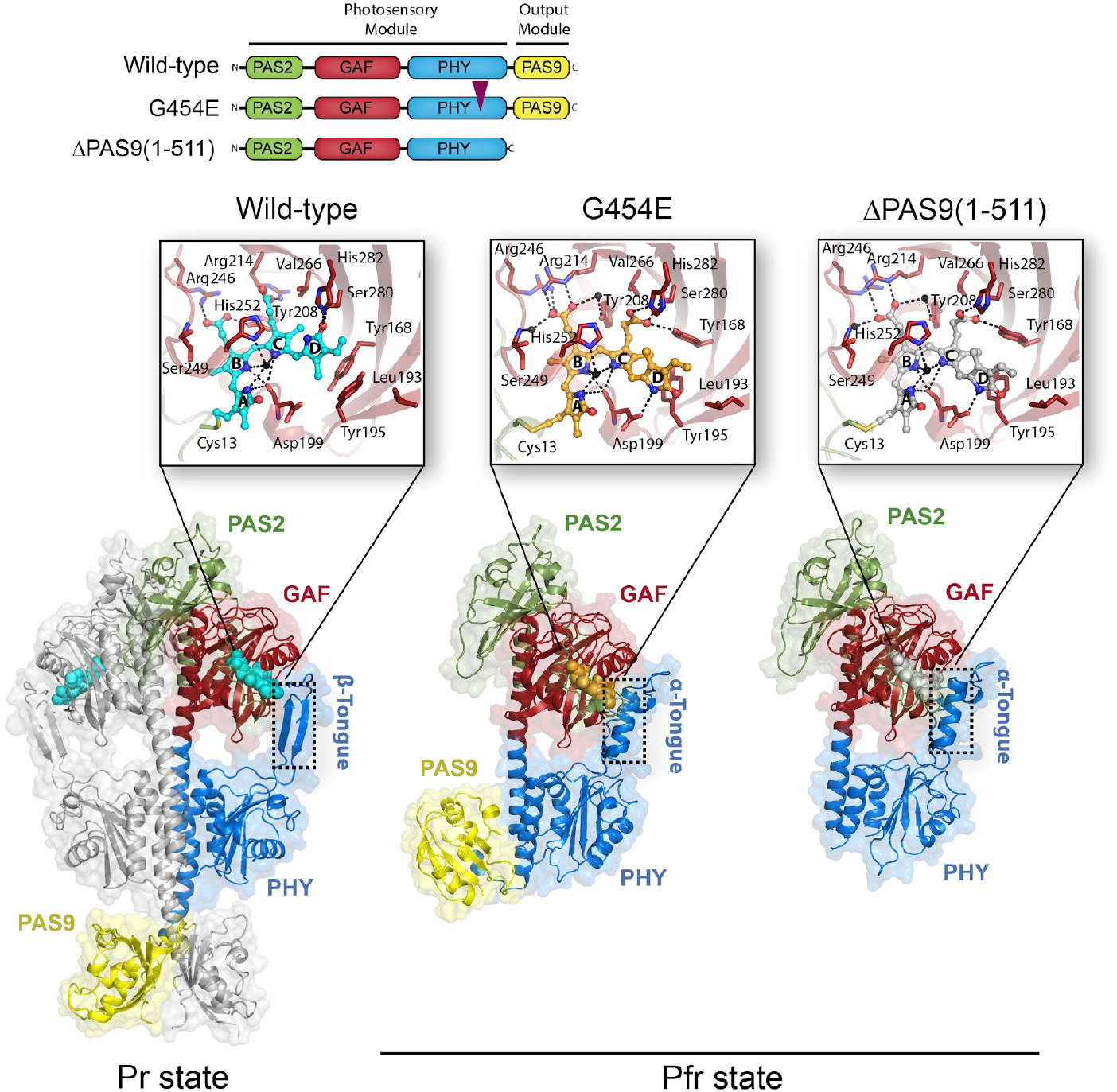
Crystal structures of *Xcc*BphP variants in the Pr and Pfr states. The domain architecture of the wild-type, G454E and ΔPAS9(1-511) variants is schematized showing the PAS2 (green), GAF (red), PHY (blue), and PAS9 (yellow) domains; the same color code is applied to all figures in this work. The domain composition of the photosensory module (PSM) and output module (OM) are indicated. A purple triangle indicates the position of the single amino acid change within the G454E variant. Ribbon representations of the different *Xc*cBphP variants: wild-type (Pr state), G454E (Pfr state), and ΔPAS9(1-511) (Pfr state) as defined in their respective asymmetric units. The tongue regions are highlighted in dashed boxes, exposing a two-stranded antiparallel β-sheet conformation in the Pr state, and an α-helical conformation in the Pfr state. The different domains are labelled and the solvent-accessible surface calculated by PyMOL is shown in the background. *Insets*: detailed views of the chromophore-binding pocket from each structure. The BV chromophore is covalently bound to Cys13 (from the PAS2 domain) and buried into the GAF domain. BV is shown as spheres with carbon atoms in cyan (wild-type), orange (G454E), or gray [ΔPAS9(1-511)], oxygen atoms in red, and nitrogen atoms in blue. The four BV pyrrolic rings are indicated as A, B, C and D. The most relevant residues are depicted as sticks. Structural water molecules (including the pyrrole water molecule coordinated by the NH groups of rings A, B and C, and the Asp199 carbonyl group) are represented as black spheres. Important polar interactions are shown as dashed lines.

### The Pr and Pfr overall structural features

The wild-type full-length protein bearing the PSM and the complete C-terminal OM crystallized in a characteristic Pr state evidenced by (i) the *ZZZssa* chromophore configuration, (ii) the side chain conformation of the conserved Tyr168-Tyr195 pair that surrounds ring D, and (iii) the two-stranded antiparallel β-sheet tongue conformation (**Figure 2**). The Pr crystal structure obtained in this work is essentially equivalent to the former wild-type protein structure (*31*), revealing a head-to-head parallel dimer with C_α_-r.m.s.d. values of 0.57 and 0.56 Å for chains A (590 aligned residues) and B (599 aligned residues), respectively. Moreover, no significant structural differences are perceived in the BV interaction pocket. However, the electron density in ring D from the BV chromophore is clearly defined, allowing a univocal determination of the chromophore in the *ZZZssa* configuration and therefore unambiguously validating the Pr state (**Figure 2, *inset* and Figures S5 and S6**).

The structures of the full-length G454E (PSM and the complete C-terminal OM) and ΔPAS9(1-511) (only PSM) proteins are congruent with other PSMs from BphPs solved in the Pfr state (*12*, *22*, *24*, *28*, *33*, *35*) revealed by (i) the *ZZEssa* chromophore configuration, (ii) the side chain conformation of the conserved Tyr168–Tyr195 pair surrounding ring D, and (iii) the α-helical tongue conformation, among other Pfr structural hallmark features (**Figure 2 and Figure S7**). Remarkably, the OM (PAS9 domain) from G454E is found in a contracted position with respect to the PSM as a result of a conspicuous break in the PHY-OM helical linker (**Figure 2**). This key structural feature will be developed in further sections.

Interestingly, both proteins show highly similar PAS, GAF, and PHY domain structures, revealed by a pairwise alignment between their main-chain C_α_ atoms yielding r.m.s.d. values of 0.41 Å (113 aligned residues), 0.39 Å (166 aligned residues), and 1.08 Å (164 aligned residues), respectively (**Figure S8**, ***upper panel***). Furthermore, the conserved figure-of-eight knot that encloses the chromophore pocket remains unaltered between both structures **(Figure S9)**. However, a global comparison shows that the most prominent differences between both Pfr structures lie in the position of their PHY domains, which display a slight translational displacement of ~3 Å between each other, along with a subtle change on the trajectories of the GAF-PHY and PHY-OM helical linkers (**Figure 3A**). These structural differences, possibly inflicted by the presence / absence of the PAS9 domain (**Figure 3A**, ***left inset***), do not alter the tongue conformation between both proteins (**Figure 3A**, ***right inset***). No changes are observed around the position 454, which is located in the tongue loop, demonstrating that the G454E substitution does not affect the tongue overall structure. Moreover, both the residue positions as well as the bilin ligand location on the chromophore binding pocket are identical in the two structures, showing the expected interactions for the Pfr state around the BV molecule (see details in the next section) (**Figure 2, insets and Figures S5, S6 and S7**).

**Figure 3.**
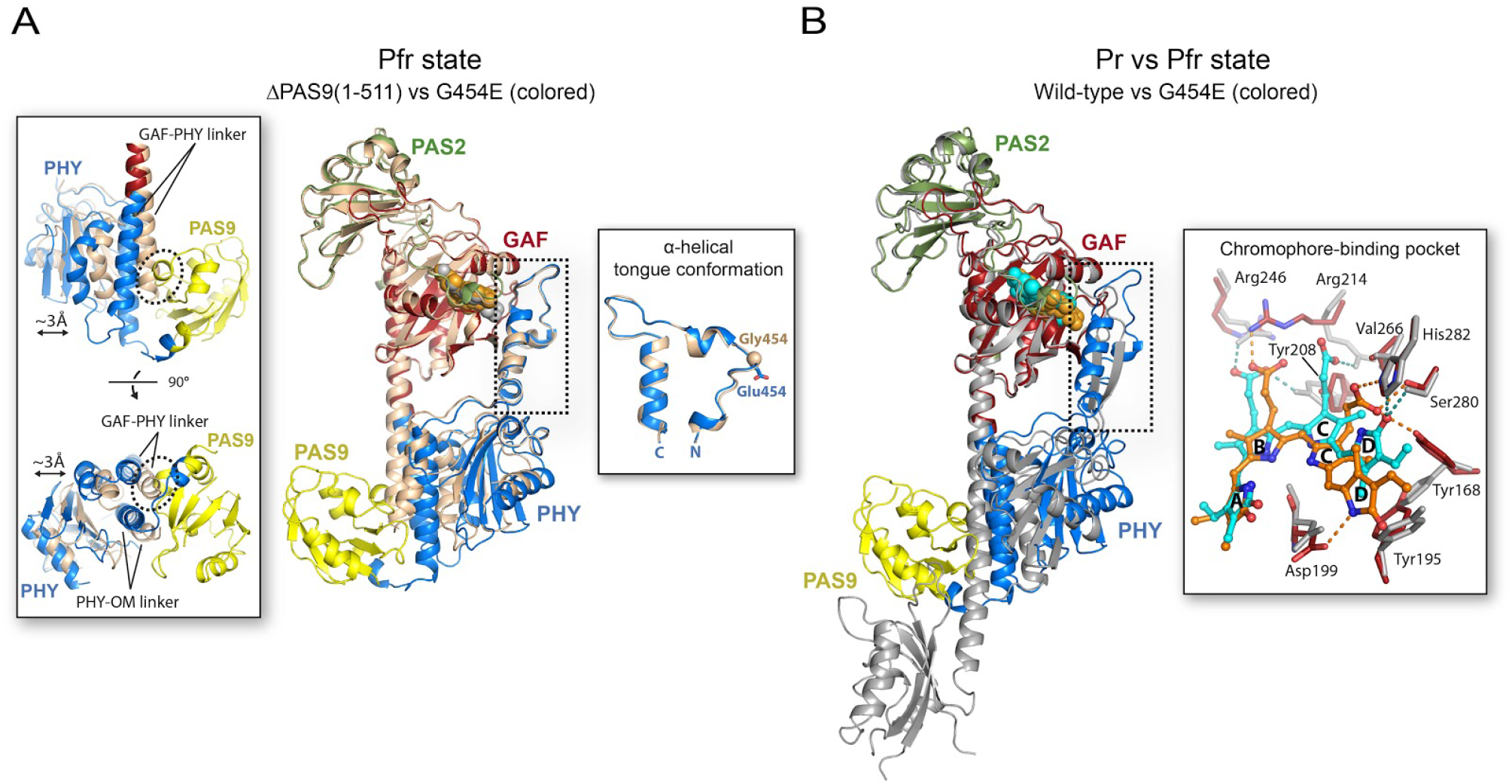
Global comparison of the tertiary structure of *Xcc*BphP variants in the Pr and Pfr states. A) Structural alignment between the Pfr structures of ΔPAS9(1-511) (wheat) and G454E. The tongue region is boxed in a dashed rectangle. The BV molecules are represented as spheres, colored in gray for ΔPAS9(1-511) and orange for G454E. The different domains are labelled. *Left inset*: the most prominent differences between both structures are shown in two different orientations rotated by 90°. The displacement of ~3 Å between the position of the PHY domains and the changes in the trajectories of the GAF-PHY and PHY-OM helical linkers are indicated. The potential clashes between the GAF-PHY helical linker, as defined in ΔPAS9(1-511) structure, and the PAS9 domain, as defined in G454E structure, are highlighted by dashed ovals. *Right inset*: α-helical tongue conformation from both proteins. The view is rotated 90° with respect to the main panel. Gly454 from ΔPAS9(1-511) and Glu454 from G454E are depicted as a sphere and sticks, respectively. N- and C-termini are indicated. B) Structural alignment between the full-length wild-type Pr structure (gray) and the full-length G454E Pfr structure variants (domains colored according to **Figure 2**). The tongue region is boxed in a dashed rectangle. The BV chromophores are represented as spheres, colored in cyan (wild-type) or orange (G454E). The different domains are labelled. *Inset*: structural differences on the chromophore-binding pocket from both proteins. BV is shown as capped sticks with carbon atoms in cyan (wild-type) or orange (G454E), oxygen atoms in red, and nitrogen atoms in blue. The four BV pyrrolic rings are indicated as A, B, C and D. The most relevant residues are depicted as sticks; the color code for G454E is carbon atoms in red within the GAF domain, oxygen atoms in red, and nitrogen atoms in blue; wild-type residues are grey-colored. Important polar interactions are shown as dashed lines colored in cyan (wild-type) or orange (G454E). Structural alignments in (A) and (B) were performed on the GAF domain.

A structural contrast between the Pr and Pfr structures reveals notable changes in different regions of the protein as illustrated in **Figure 3B**. For clarity purposes, we will describe them in detail in the following sections: (i) Conformational differences in the chromophore-binding pocket, (ii) Structural rearrangements around the chromophore-binding pocket and the tongue interconversion, (iii) Large-scale conformational changes in the helical spine and the OM, and (iv and v) the Pr and Pfr quaternary arrangements.

### Conformational differences in the chromophore-binding pocket

In the Pr and Pfr structures, the BV chromophore is embedded in a cleft inside the GAF domain and covalently bound by a thioether linkage between the C3^2^ atom of ring A and a conserved cysteine residue (Cys13) in the N-terminal extension of the PAS2 domain (**Figure 2**, ***insets* and Figure S6**). The global folding of the PAS2 and GAF domains, including the figure-of-eight knot, remain unaltered between both photostates (**Figure 3B, Figures S8**, ***bottom panel* and S9**). However, conformational changes into the chromophore are recognized, which give rise to spatial differences on the enclosed side-chain residues and, consequently, rearrangements of the interaction network.

As noted previously, the Pr state presents a *ZZZssa* chromophore configuration, whereas the transition to the Pfr state involves an isomerization of the ligand with a ~180° rotation in the ring D showing a *ZZEssa* configuration (**Figure 2**, ***insets* and Figure 3B, *inset***). In concordance with other reported BphP structures (*8*), this configurational motion is accompanied by a synchronized flip-and-rotate movement of two highly conserved aromatic residues, Tyr168 and Tyr195, located next to ring D, along with a slight rearrangement of the Tyr255 side chain (**Figures 3B, *inset* and 4A**). This local reorganization is associated with a series of structural motions outside the chromophore-binding pocket explained in the next section. Moreover, the hydrogen bonds formed among the ring D carbonyl oxygen and the side chains of Ser280 and His282 in the Pr state are replaced in the Pfr structures by a new hydrogen bond between the NH group from ring D and the Asp199 carboxylate, which is part of the highly conserved DIP motif (residues 199-201), (**Figures 2, *insets*, 3B, *inset* and Figure S6**). Accordingly, it has been reported in other BphPs such as Agp1 from *Agrobacterium fabrum* (also known as *A. tumefaciens*), *Dr*BphP from *Deinococcus radiodurans* and *Pa*BphP from *Pseudomonas aeruginosa*, that the abolition of the aspartate residue in this motif typically blocks the protein photocycle in an intermediate state that disrupts the Pr-to-Pfr progression (*18*, *36*, *37*).

A network of interactions among the NH groups of rings A, B and C, the Asp199 carbonyl group, the conserved pyrrole water molecule and the N^δ^ atom of His252 are observed both in the Pr and the Pfr states (**Figure 2, *insets* and Figure S6**). However, the positions of BV rings B and C (but not A) are clearly displaced between both photostates, altering the hydrogen bond network of their propionate groups (**Figure 3B, *inset***). In the Pr state, the two BV propionates are within hydrogen-bonding distance from the side chains of Tyr208, Arg214, and Arg246; however, the chromophore displacement observed in the Pfr state is associated with a different interaction network (**Figures 2, *insets*, 3B, *inset* and Figure S6**). In the Pfr state, the propionate group of ring B is directly stabilized by the side chain of Arg214, and indirectly by Tyr208, Arg246 and Ser249 *via* ordered water molecules. Furthermore, the propionate group of ring C is relocated into a new hydrogen-bonding environment constituted by the side chains of Tyr168 (from the tyrosines flip-and-rotate mechanism), Ser280 and His282 (**Figures 2, *insets*, 3B, *inset* and Figure S6)**.

In order to assess the contribution of relevant residues for Pr and Pfr stability, as mentioned above and in the following sections, we designed a set of mutations and evaluated their photochemical behaviour, using the dark conversion rate (half-life) and the Pr:Pfr proportions at thermal equilibrium as proxies for stability **(Figure 1, Table 1 and Figures S2-S4)**. For the sake of simplicity in the photochemical classification of the variants, we define the types as “bathy”, “bathy-like”, “canonical-like” and “canonical” when they exhibit a Pfr enrichment in thermal equilibrium above 90%, between 50 and 90%, between 10 and 50% and below 10%, respectively (**Figure 1**).

To validate the displacement of the propionate group of ring C observed between both photostates, we designed two substitutions that exchange the polarity and hydrophobicity in two close residues, Val266 by serine (V266S) and Ser280 by valine (S280V) (**Figures 2, *insets*, and 3B, *inset***). Both substitutions make a significant change in the preferred state in equilibrium, turning the *Xcc*BphP into canonical-like and canonical types, respectively (**Figure 1 and Table 1**). As observed in the structures, the Val266 side chain is near the ring C propionate in either the Pr or Pfr conformations. However, the extra hydrogen-bond introduced by the serine hydroxyl group in V266S seems to stabilize this propionate group from BV in the *ZZZssa* configuration. In contrast, the elimination of the serine hydroxyl group in S280V may destabilize this group from BV in the *ZZEssa* configuration, impairing the Pfr state.

### Structural rearrangements around the chromophore binding pocket and the tongue interconversion

In *Xcc*BphP, the tongue is constituted by residues 446-483, which are organized in the highly conserved motifs W(G/A)G (452-454), PRxSF (471-475), and (W/Y/F)x(E/Q) (478-480) (*31*).

In the Pr state, the tongue adopts a β-hairpin fold and the Trp452 residue is located in the entrance β-strand within the tongue-GAF interface, while Trp478 is situated in the exit β-strand and exposed to the protein surface (**Figure 4A, *left panel***). Furthermore, Ala453, Gly454, Pro471, Arg472 and Phe475 are positioned in the β-hairpin loop adjacent to the chromophore-binding pocket, and stabilized mainly by hydrogen bonds among the side chains of Arg472, Asp199, and Tyr255 (**Figure 4A, *left panel***). Remarkably, residues 456-470 from the β-hairpin loop were not visible into the electron density map. This distal portion of the tongue is not resolved in several other BphP Pr structures either, indicating a high flexibility for this region.

**Figure 4.**
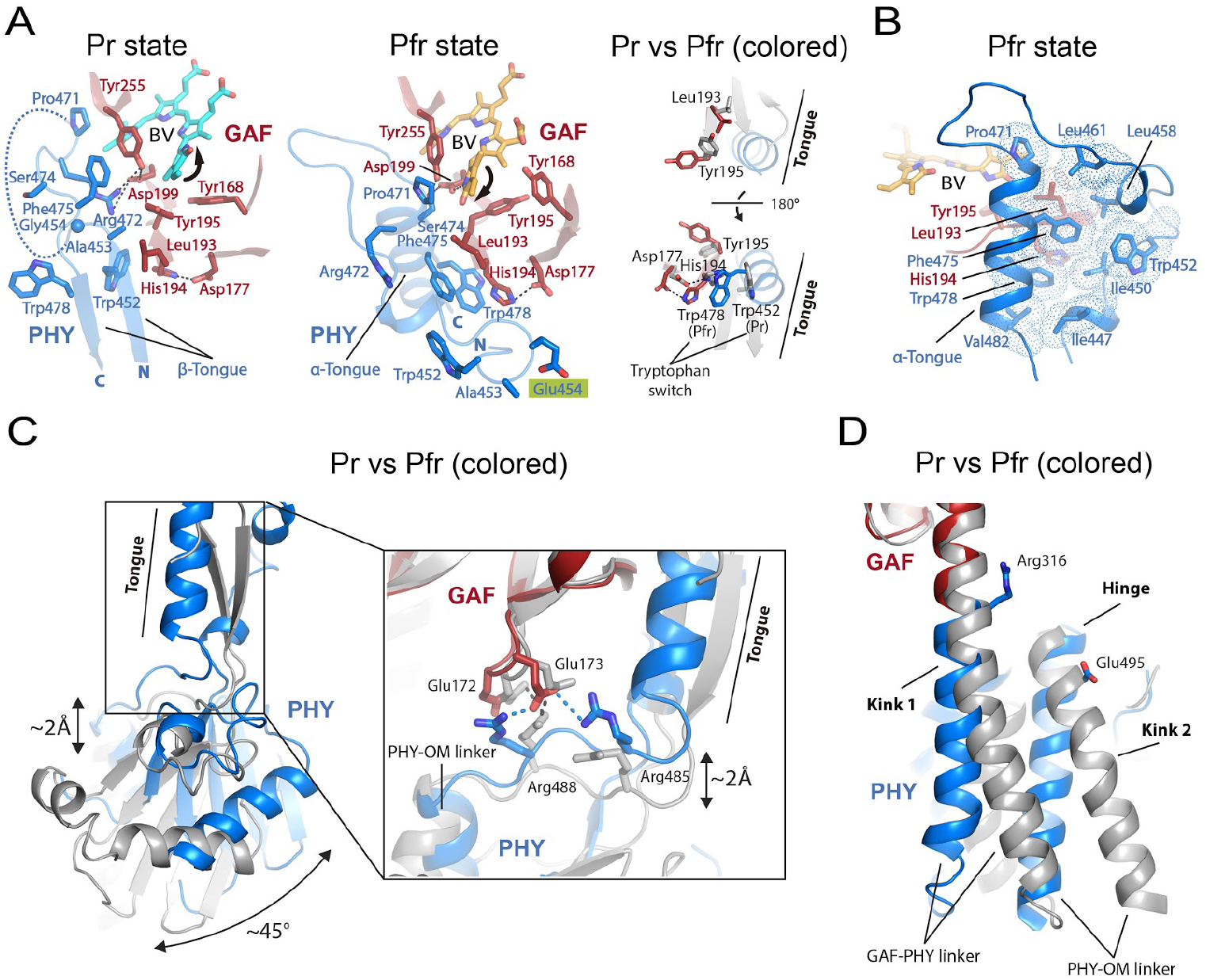
Structural rearrangements around the chromophore-binding pocket and the tongue interconversion between the full-length Pr and Pfr structures. A) *Left and central panels*: detailed view of the main interactions observed in the Pr structure of wild-type *Xcc*BphP and the Pfr structure of G454E, respectively. The most relevant residues are depicted as sticks, and colored following the domain color code of **Figure 2**. Gly454 from the wild-type Pr structure and Glu454 (highlighted in a green box) from the G454E Pfr structure are depicted as a sphere and sticks, respectively. Important polar interactions are shown as dashed lines. The disordered region of the β-hairpin loop in the Pr structure is represented by a curved dashed line. BV is shown as sticks with carbon atoms in cyan (wild-type) and orange (G454E), oxygen atoms in red, and nitrogen atoms in blue. The BV ring D movements upon photoconversion are represented with black arrows. The GAF and PHY domains, the N- and C-termini, and the tongue region are indicated in both panels. *Right panel*: structural contrast between both structures (Pr in gray and Pfr colored following **Figure 2**) displayed in two different orientations rotated by 180°, emphasizing the interactions by Leu193 and His194. Tryptophan residues from the Trp switch and the tongue region are indicated. B) Hydrophobic pocket in the tongue-GAF interface of the Pfr state. The most relevant residues and the BV are shown as sticks and colored according to the central panel in (A). The hydrophobic contacts are represented by a cloud of dots. C) The tongue fold interconversion between the β-sheet and α-helix conformations and the PHY domain repositioning. The Pr (gray) and Pfr structures (colored according to **Figure 2**) are aligned on the GAF domain. The displacement of ~2 Å between the GAF and PHY domains as well as the reorientation of the PHY domain of ~45° are indicated. *Inset*: detailed view of the changes in the salt-bridge clusters from the interfacing GAF/PHY network. Interactions are shown as dashed lines colored in grey (wild-type *Xcc*BphP) or blue (G454E). The GAF and PHY domains, and the PHY-OM helical linker are indicated. D) Changes on the helical spine. The Pr (gray) and Pfr (colored according to **Figure 2**) structures are aligned on the GAF domain. The bending regions in the GAF-PHY and PHY-OM helical linkers from Pr structure are denoted as “Kink 1” and “Kink 2”, respectively. The pivot point in the trajectory of the PHY-OM helical linker is indicated as “Hinge” at the beginning of the helix. The residues Arg316 (GAF domain) and Glu495 (PHY domain) involved in the Pr dimer interface (see **Figure 6A, *upper panel***) are shown as references.

On the contrary, in the Pfr state the tongue undergoes a conformational change interconverting the entrance and exit β-strands in a random loop and an α-helix, respectively (**Figures 3B and 4A, *central panel***). As a result, an exchange of the conserved residues Trp452/Trp478 in the tongue-GAF interface takes place in concordance with the Trp switch model previously proposed (*29*, *38*). Concordantly, the Trp478 residue located in the tongue α-helix is now placed within the tongue-GAF interface, and Trp452 is exposed in the tongue loop facing the solvent along with Ala453 and the variant residue Glu454 (**Figure 4A, *central panel***). In addition, Pro471, Ser474 and Phe475, which are settled also in the α-helix of the tongue, close tighter the tongue-chromophore pocket interface by setting new interactions: (i) Pro471 contacts the vinyl side group of the BV ring D and Tyr255, (ii) Ser474 is located at van der Waals distance from the Asp199 and Tyr255 side chains (**Figure 4A, *central panel***), and (iii) Phe475 is associated to a predominantly hydrophobic pocket constituted by Leu193 and the tongue residues Ile447, Ile450, Trp452, Leu458, Leu461, Pro471, Trp478, and Val482 (**Figure 4B**).

As already stated, the structural connection between the bilin photoisomerization and the tongue interconversion is not completely understood. The crystal structures reported here indicate that upon BV photoisomerization (ring D rotation), the phenolic side chains of the tyrosine triad 168-195-255 in the immediate surroundings of ring D change their positions. The Tyr195 residue lies at a β-strand along with its preceding residues Leu193 and His194, which participate in the tongue-GAF interface (**Figure 4A**). Due to a geometrical feature of the β-strand structure, the side chains of Leu193 and Tyr195 are inside the chromophore-binding pocket, whereas the side chain of His194 is located outside. This structural organization places the side chains of Leu193 and Tyr195 in close distance. Accordingly, in the Pr state the OH group of Tyr195 seems to exclude the Leu193 aliphatic side chain from the chromophore binding domain by steric effects (**Figure 4A, *left and right panels***), while in the Pfr state its location allows Leu193 to position its side chain in that area next to ring D (**Figure 4A, *central and right panels***). As a result, the distance between the Leu193 side chain and the BV ring D decreases considerably from ~7 in Pr to ~3 Å in Pfr. This movement in the Leu193 side chain seems to affect the hydrophobic interactions observed in the tongue-GAF interface of the Pfr state (**Figure 4B**). Consequently, the side chain of His194 flips in the opposite direction (probably by a subtle main chain torsion due to the movement of the adjacent residues Leu193 and Tyr195) and pulls the Asp177 residue by a hydrogen bond (**Figure 4A**). Strikingly, the His194 motion is accompanied by the Trp switch model described above, through a cation-π or π-π stacking interaction between its imidazole ring and the indole ring of Trp452 in the Pr state or Trp478 in the Pfr state. Moreover, the Tyr255 movement allows or hinders the Phe475 side chain positioning near the BV molecule in the Pr and Pfr states, respectively.

To challenge these observations and to gain insight into their roles in the Pr-Pfr photocycle, we designed another set of mutations affecting some key residues. The replacement of His194 by Ala (H194A) presents a ~2.7-fold increase of the Pr proportion at equilibrium indicating that abolition of the cationic imidazole ring affects more the Pfr state stabilization (**Figure 1 and Table 1**). In this line, the contacting residue Asp177 does not seem to play a significant role in the His194 flipping, as revealed in the D177A variant, whose photochemical behaviour is very similar to the wild-type protein. Additionally, W478A dramatically shifts the Pr:Pfr ratio at equilibrium to 9:1, turning it into a canonical-like type. This evidence supports the notion that there is a functional connection between His194 and Trp478 (linking the chromophore binding pocket and the tongue), which is crucial for the Pfr state. Similarly, the Ser474 by Glu substitution (S474E) converts the phytochrome into a canonical type, driving the dark equilibrium towards an almost full conversion to the Pr state and with a surprisingly low half-life (60 min, **Figure 1 and Table 1**). In concordance with the Pfr structure, the insertion of a carboxylate group in this position might provoke a repulsive electrostatic effect with the Asp199 side chain, validating the Ser474 role in the tongue-GAF interface in this photostate (**Figure 4A, *central panel***).

Interestingly, the Trp452 by Ala substitution (W452A) exhibits a slight change in the Pr:Pfr ratio at equilibrium, indicating that Trp452 does not play a significant role in the stabilization of the Pr state. Nevertheless, the significant increase of the Pr-to-Pfr dark conversion rate (3.4 fold) relative to the wild-type suggests that it may be participating in the stabilization of a tongue intermediate state. It is noteworthy that substitutions on nearby residues (*i.e.* A453E and G454E) transform the phytochrome into a typical bathy type, where an almost complete Pr-to-Pfr dark conversion occurs and at significantly accelerated rates compared to the wild-type with ~24- and ~15-fold increases, respectively (**Figure 1 and Table 1**). These results are in total agreement with the structural data, where both residues are involved in GAF-PHY interface in the Pr state, but not in the Pfr (**Figure 4A, *left* and *central panels***). Therefore, the incorporation of a negative charge in both positions may dislocate the GAF-PHY interface observed in the Pr state, thus favouring the Pfr state. Regarding Gly454, it is likely that the high conformational flexibility of this residue due to its sidechain-less backbone is necessary for Pr stabilization. This scenario is supported by a recent work from our group in which we showed that substituting Gly454 by almost any other amino acid resulted in Pfr-favoured variants (*34*).

### Large-scale conformational changes in the helical spine and the OM

As reported previously (*22*), the tongue fold interconversion between the β-sheet and α-helix conformations implies an elongation or a shortening of this peptide segment, respectively. As a consequence, in *Xcc*BphP a push (Pr) / pull (Pfr) displacement of around 2 Å between the GAF and PHY domains takes place, along with a prominent reorientation of the PHY domain of ~45°, which alter the GAF/PHY interface between both photostates (**Figures 3B and 4C**). In the Pr state, a network of salt-bridges involving residues Glu172 and Glu173 (from the GAF domain), and Arg488 (from PHY) are observed, while in the Pfr state the interfacing GAF/PHY network includes the residues Glu173 (from GAF), and Arg485 and Arg488 (from PHY) (**Figure 4C, *inset***). The PHY domain repositioning does not affect the topology of the secondary structure elements from its globular part, although local changes can be observed on their 3D arrangement (**Figure S8, *bottom panel***), as detailed below.

In line with the PHY domain rearrangement, changes on the helical spine are noted. In the full-length Pfr structure the GAF-PHY helical linker is fairly straight, while in the Pr structure it is slightly kinked (kink 1) (**Figures 3B and 4D**). Moreover, a pivot point at the beginning of the PHY-OM helical linker (hinge) is observed, which affects the helix trajectory (**Figures 3B and 4D**). As the Pr head-to-head dimer interface mainly lies on the GAF-PHY and PHY-OM helical linkers (see **Figure 6A, *upper panel***), their structural rearrangements affect the positions of the interfacing residues, and consequently the dimeric interactions. Among them, it is worth noting the Glu328/Arg501’ and Glu328’/Arg501 salt-bridges, which are found anchoring both helical linkers in the Pr dimer. Remarkably, on this photostate, Arg501 is arranged in the PHY-OM helical linker where another kink is noted (kink 2, **Figure 5A)**. Conversely, in the Pfr state, Arg501 is settled on a straight α-helix forming a salt-bridge with Glu505 from the same protomer.

**Figure 5.**
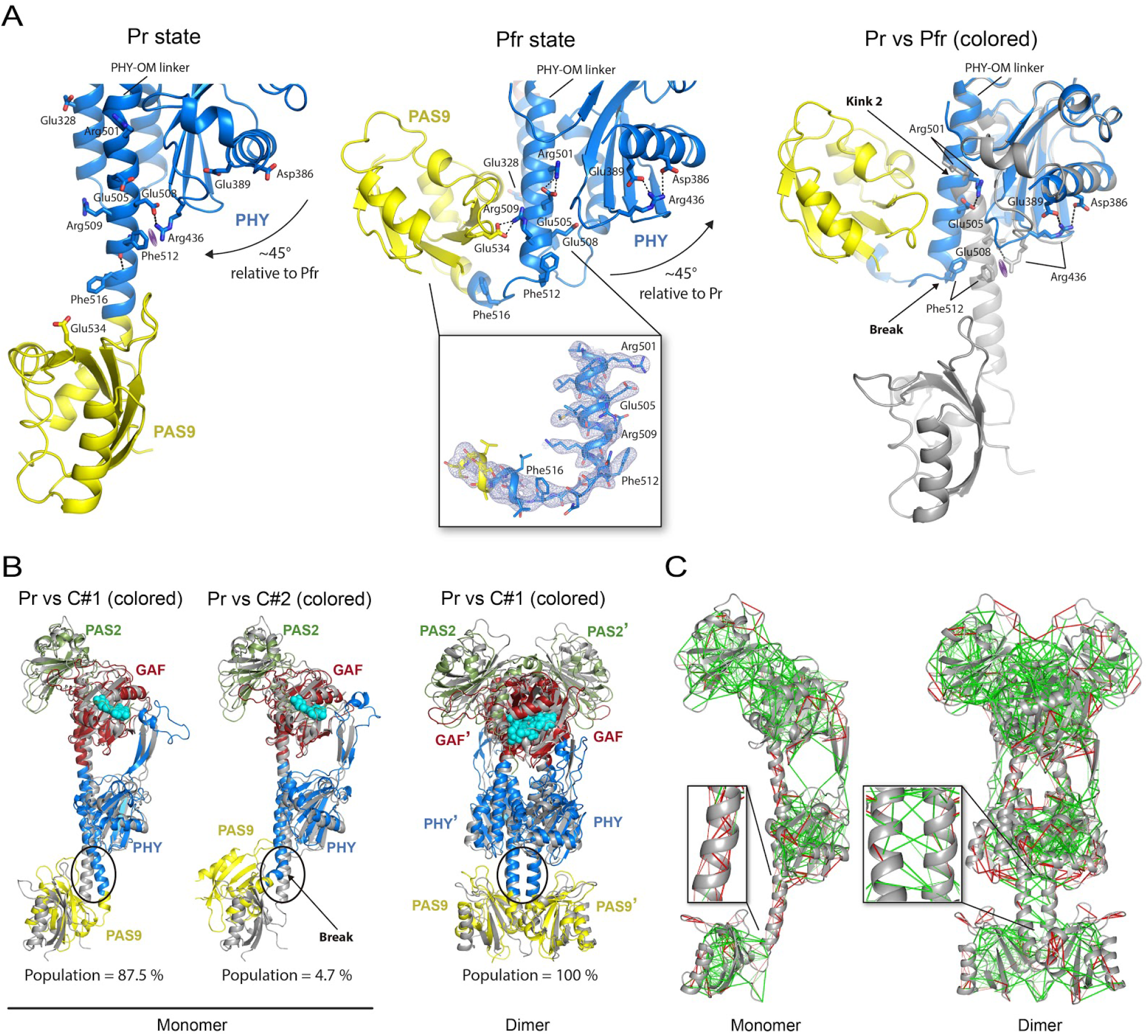
Large-scale conformational changes in the PHY-OM helical linker and the OMs between the full-length Pr and Pfr structures. A) *Left and central panels:* detailed view of the main interactions observed in the wild-type *Xcc*BphP Pr structure and the in the G454E Pfr structure, respectively. The most relevant residues are depicted as sticks, and colored following **Figure 2** domain color code with oxygen atoms in red, and nitrogen atoms in blue. The PHY domain repositioning between Pr and Pfr (~45°) is indicated with arrows in both panels. Important polar interactions are shown as dashed lines. The cation-π interaction between Arg436 and Phe512 is represented by a purple meshed oval. The PHY and PAS9 domains are labelled. *Inset of the central panel:* the final *2mF*_o_–*DF*_c_ electron density map (light blue mesh) contoured at the 1.0 σ level is exhibited defining the break at the central region of the PHY-OM helical linker in the Pfr structure. *Right panel:* contrast between both structures. The Pr (gray) and Pfr (colored according to **Figure 2**) structures are aligned on the PHY domain. The bending (Pr) and the disruption (Pfr) in the PHY-OM helical linker are denoted as “Kink 2” and “Break”, respectively. The most relevant polar interactions from each structure are shown as in the left and central panels. B) Structural alignment between the Pr crystal structure (gray) and MD averaged models (colored according to **Figure 2**, Pr structure). *Left panel:* MD clusters C#1 and C#2 obtained from a monomer of the Pr structure of wild-type *Xcc*BphP. *Right panel:* MD cluster C#1 obtained from the dimeric Pr structure of wild-type *Xcc*BphP with chains A and B (indicated as prime). The population fraction (%) of each cluster is indicated in both panels. The region around the “break” observed in the PHY-OM helical linker of the G454E Pfr structure is stressed with an oval. The different domains are labelled. C) Local frustration patterns on the monomeric (left) and dimeric (right) Pr scaffolds using the Frustratometer server (49). The protein backbone is displayed as gray ribbons, and the inter residue interactions with solid lines. Minimally frustrated interactions are shown in green, highly frustrated interactions in red, and neutral contacts are not drawn. *Insets*: Expanded views of the interactions determined around the “break” region observed in the PHY-OM helical linker of the G454E Pfr structure.

**Figure 6.**
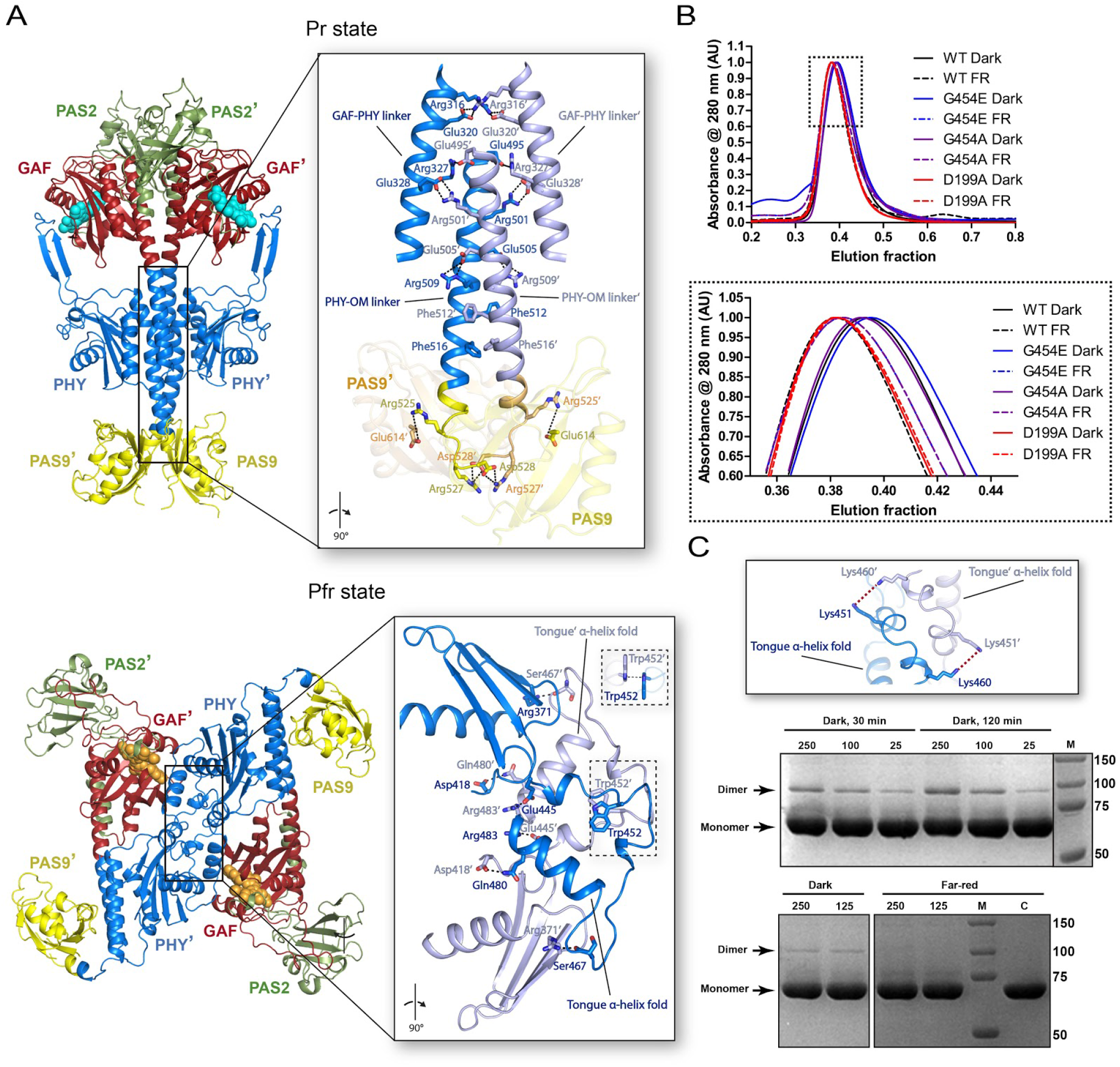
Quaternary arrangements of the full-length Pr and Pfr structures. A) *Upper panel*: Head-to-head dimer assembly of the wild-type Pr crystal structure as found in its asymmetric unit. *Bottom panel*: Head-to-tail crystallographic dimer assembly of the G454E Pfr crystal structure constructed by means of the *−x, −y, z* symmetry operator of the *I*4_1_ space group. Structures are shown in ribbon representation, and colored following **Figure 2** domain color code with the two protomers depicted in different shades. *Insets*: Detailed view (rotated 90° with respect to main panels) of the respective dimerization interfaces encompassing the GAF-PHY and PHY-OM helical linkers from the helical-spine in the Pr structure, and the tongue region in the Pfr structure. The most relevant interactions are depicted as sticks, and colored according to the chain with oxygen atoms in red, and nitrogen atoms in blue. Important polar interactions are shown as black dashed lines. Two views of the π-π stacking interaction between Trp452/Trp452’ (from the Trp switch) in the Pfr structure are enclosed by dashed boxes. B) SEC of dark adapted (solid lines) or far-red irradiated (FR, dashed lines) of wild-type *Xcc*BphP, G454E, G454A and D199A holoprotein variants. Dark-adapted holoproteins were subjected to a gel-filtration in a covered column, while far-red irradiated holoproteins (FR) were illuminated prior (20 min) and during the run. The entire runs are displayed in the upper panel, while the lower panel details their peaks region. Representative plots from three replicates are depicted. C) Dimer cross-linking of dark adapted or far-red irradiated wild-type *Xcc*BphP. The dark-adapted or far-red illuminated (20 min) holoproteins were incubated with the cross-linker BS3 at a 25, 100, 125 or 250 molar excess. *Upper panel*: Representation of the potential interactions (red dashed lines) between the only pair of primary amines (here Lys residues) located at cross-linking distance in the dimeric interface of the Pfr G454E crystal structure colored following A) domain color code. *Middle and bottom panels:* two SDS-PAGE gels from representative experiments. (M: markers, with MW values in kDa indicated on the right). The C lane (control) corresponds to the dark-adapted wild-type protein incubated in the absence of the cross-linker. The light conditions, incubation times and molar excess are indicated above the gel. The position corresponding to cross-linked dimer and monomer bands are indicated on the left.

To probe the relevance of the Glu328/Arg501 interactions in the photochemical mechanism, we eliminated the salt-bridges by replacing Arg501 by Ala (R501A) and evaluated the photochemical properties of this variant. Although slight changes were observed regarding the Pr:Pfr relative abundances at dark equilibrium, we observed a drastic change of 3.5-fold reduction in the Pr-to-Pfr conversion rate compared to the wild-type (**Figure 1 and Table 1**). Hence, the abolition of the Glu328/Arg501 contacts could be leading to a faster Pr-to-Pfr transition by generating a less stabilized PHY-OM helical linker kink in the Pr state.

Furthermore, an impressive break at the central region of the PHY-OM helical linker is noted in the Pfr state (hereafter referred as “break”), which places the PAS9 domain into a contracted position in comparison with the Pr state, but preserving its global folding (**Figures 2, 3B and 5A**). This unprecedented helical disruption, unambiguously defined in the electron density map (**Figure 5A and Figure S10**), turns the PHY-OM helical linker and the PAS9 domain closer in space resembling a C-like protein global folding (**Figure 3B**). Remarkably, the electron density for the PAS9 domain in this position is slightly less defined with relatively higher *B*-factor values (**Figure S11**), which denotes higher conformational flexibility favored by lacking intermolecular contacts within the crystal packing (**Figure S12**). Additionally, molecular dynamics (MD) simulations conducted on the G454E crystal structure revealed four major clusters that encompass minor structural changes with respect to the crystallographic structure (C_α_-r.m.s.d. values ranging from 1.78 to 2.10 Å, **Figure S13**) suggesting that the PAS9 domain position defined in the Pfr state is conserved under no confinement conditions as those present in the crystal.

A salt-bridge between Arg509 and Glu534 is found stabilizing the contracted PAS9 domain with the PHY-OM helical linker in the Pfr state. Interestingly, Arg509 is part of the Pr dimer interface by interacting with Glu505, the counterpart of Arg501 in Pfr, as described above (**Figure 6A)**. Therefore, there is a salt-bridge switch between Glu505 with Arg501 and Arg509 (from the neighboring protomer) in the Pfr and Pr states, respectively. Consequently, when the Pr dimer is assembled, Arg501 and Arg509 are trapped in the dimer interface, but in the Pfr state both are free to be contacted by Glu505 and Glu534, respectively.

The breaking point for the PHY-OM helical linker is located at the Phe512 residue (**Figure 5A, *central panel* and Figure S10, *inset***). In the Pr state, the phenyl group of Phe512 is within cation-π interaction distance with the cationic guanidine moiety of Arg436 from the PHY domain, which is in turn stabilized by a salt-bridge contact with Glu508 (**Figure 5A, *left panel***). Interestingly, Phe512 is found buried at the dimer interface close to Phe516 from the neighbouring chain (see **Figure 6A**). Following the mutational strategy, we aimed to simulate the PHY-OM helical linker break by introducing a proline residue at position 512 (F512P). The F512P variant photochemistry did not show a marked change of the Pr and Pfr relative stabilities at equilibrium compared to the wild-type **(Figure 1 and Table 1**), although the kinetics of its Pr-to-Pfr conversion did show a ~4-fold decrease, which is indicative of an alteration of one or more transition states. Additionally, the MW determination of the F512P apoprotein is consistent with a dimer configuration (**Table 1 and Figure S16**). Therefore, a helical disruption at this position still allows for a photoswitchable *Xcc*BphP phytochrome, with an unaltered dimeric arrangement.

In the Pfr state, the pulled PHY domain moves Arg436 ~18 Å away from Phe512, relocating it at salt-bridge distances with the intra-domain residues Asp386 and Glu389 (**Figure 5A, *central panel***). Noteworthy, in the Pr state the Phe512 carbonyl group interacts with the Phe516 (*i* + 4) amide group in a turn from the PHY-OM helical linker, while in Pfr this interaction is absent as a result of the breaking (**Figure 5A**). According to the mutational analysis, the abolition of the Phe512 phenyl group (F512A) accelerates the Pr-to-Pfr dark conversion (~2.4-fold) without changing drastically the Pr:Pfr equilibrium **(Figure 1 and Table 1**), in a similar fashion to the F512P variant, also suggesting an alteration of one or more transition states. Thus, the cation-π interaction between Arg436 and Phe512 might not play a major role in Pr stability, leaving this role potentially to the nearby salt-bridged contact Arg436-Glu508 (**Figure 5A**), but it would be involved in the photoconversion pathway.

In the same line, MD simulations on the monomeric Pr structure resulted in two clusters, one major showing a bend in the PHY-OM helical linker, and another exhibiting a break in a similar region to that observed in the Pfr full-length structure (**Figure 5B, *left panel***). Interestingly, the same computational studies performed on the dimeric Pr structure exposed no changes in this region (**Figure 5B, *right panel* and Figure S14**), suggesting that dimer dissociation is needed for the breaking to occur.

Additionally, we calculated the local frustration patterns on the monomeric and dimeric Pr scaffolds using the Frustratometer server (*39*) to analyze how the local frustration distribution in this helical region might be affected by the oligomeric state. As specific regions in protein structures require a great propensity to be frustrated to facilitate both motion and interaction with partners, these frustrated areas can be flexible or cracked. Thus, sites of high local frustration correlate with regions involved in protein function (e.g., binding sites, allosteric transitions) (*39*). Interestingly, while in the Pr monomer the breaking area observed in the G454E crystal structure is consistently found to be highly frustrated, in the dimer it is minimally frustrated **(Figure 5C, and Figure S15).**

Taken together, the computational studies suggest that the PHY-OM helical linker in this region is flexible and frustrated, and consequently prone to be kinked when it is not part of the helical dimeric interface as in the Pr state.

### Pr quaternary arrangement

As previously reported, the Pr state is assembled as a parallel head-to-head dimer (*31*). The dimeric interface (3263 Å^2^) involves 89 residues per protomer (15% of the total residues) mainly settled along the helical spine that interact by a series of salt-bridges, hydrogen bonds and non-bonded contacts from side to side at the dimer interface. Approximately, 65% of the interfacing residues are located between neighbouring GAF-PHY linkers, PHY-OM linkers and PAS9 domains that include 12 potential salt-bridged pair residues (**Figure 6A, *upper panel***).

We focused on the Arg525/Glu614’, Arg525’/Glu614, Arg527/Asp528’ and Arg527’/Asp528 salt-bridges, which stabilize the neighboring PHY-OM linkers and PAS9 domains. To assess their relevance in the Pr dimer formation, we generated a full-length variant replacing the Arg525 and Arg527 residues by Glu (opposite charge and similar side chain size), named R525E-R527E, as well as two truncated versions lacking the OM, named ΔPAS9(1-525) and ΔPAS9(1-527). Static light scattering coupled to size exclusion chromatography (SLS-SEC) experiments revealed that the three protein constructs predominantly exhibit monomeric species in solution with a minor dimeric population (**Figure S16 and Table 1**), in line with ΔPAS9(1-511), which is a monomer exclusively (*31*). This is indicative that the salt-bridges together with the hydrogen bonds and non-bonded contacts along with the neighbouring PHY-OM linkers and PAS9 domains are individually necessary but not sufficient to sustain the head-to-head dimer assembly. Analogously to the SLS-SEC data, ΔPAS9(1-525) and ΔPAS9(1-527) show a lower Pr-to-Pfr dark conversion half-lives and a higher Pr:Pfr at equilibrium compared to the wild-type version, although these changes are slightly less pronounced than in ΔPAS9(1-511) (**Figure 1 and Table 1**). R525E-R527E exhibited a considerably faster Pr-to-Pfr dark conversion rate (7.3-fold) compared to the wild-type version and an almost complete Pfr enrichment at dark equilibrium (97%) (**Figure 1 and Table 1)**. Taken together, these results indicate that the Pfr state is favoured with respect to the Pr state when the integrity of the Pr dimeric arrangement is hindered.

### Pfr quaternary arrangement

In contrast to the non-crystallographic parallel dimer assembled in the Pr state, the G454E Pfr structure shows only one polypeptide chain in the crystal asymmetric unit (**Figure 2**). However, the apoprotein G454E variant undoubtedly behaves as a dimer in solution according to SLS-SEC experiments (**Figure S16 and Table 1**). An in-depth search for symmetry-related crystallographic partners in the G454E crystal packing revealed an unusual head-to-tail quaternary assembly where both protomers are aligned nearly antiparallel and related by a two-fold symmetry axis that can be constructed by means of the −*x*, −*y*, *z* symmetry operator of the *I*41 space group (**Figure 6A, *bottom panel* and Figure S12**). This potential quaternary structure is exceptionally settled on the PHY domain from both protomers predominantly at the tongue regions sharing an interface area of 1080 Å2 (7.0% total residues of each chain). Outstandingly, the dimer interface displays an antiparallel-displaced (or offset stacked) π-π stacking interaction between Trp452/Trp452’ (from the Trp switch) along with the salt-bridge contacts Glu445/Arg483’, Glu445’/Arg483, and a series of hydrogen bonds constituted by Arg371/Ser467’, Arg371’/Ser467, Asp418/Gln480’ and Asp418’/Gln480 (**Figure 6A, *bottom panel***). Interestingly, the substitution W452A generates a formation of higher-order oligomeric arrangements by SLS-SEC measurements (**Figure S16 and Table 1**), demonstrating that Trp452 plays an important role in the *Xcc*BphP oligomerization. It is worth noting that the W478A substitution affecting the other residue involved in the Trp switch does not alter the dimeric arrangement of the apoprotein (**Figure S16 and Table 1**), in agreement with the observation that this residue is not involved in the dimer interface in either photostate.

The mutated Glu454 residue is also located at the tongue region in the dimer interface likely interacting with the protomer partner Lys460. However, no clearly defined electron density was observed around the latter residue side chain in order to unambiguously determine this interaction. Moreover, the A453E and G454A variants, which exhibit a highly similar photochemical behaviour to G454E, also show a dimeric assembly in solution **(Figure S16 and Table 1)**, suggesting that the glutamate substitution does not drive the crystallographic head-to-tail dimer but might contribute to its stabilization. Noticeably, the double mutant W452A-G454E restores the dimeric state **(Figure S16 and Table 1)**, which indicates that G454E governs the dimeric arrangement, in agreement with the fact that the G454E variant favours more the Pfr photostate than the W452A variant (**Figure 1 and Table 1**). Collectively, these findings suggest that the effects of the G454E substitution is mainly on the photochemical behavior, which is ultimately driving the dimeric head-to-tail arrangement.

Interestingly, the crystallographic dimer observed in G454E is absent in the ΔPAS9(1-511) crystal packing (**Figure S12**), which is in agreement with the monomeric behaviour observed in solution for this construct (*31*). As shown before, the PHY domain in G454E is more exposed to the solvent than in ΔPAS9(1-511), possibly pushed by the OM (**Figure 3A, *left inset***). For this reason, we hypothesize that the Pfr dimer assembly of G454E is a consequence of the PHY domain displacement described above.

The Pr head-to-head dimeric arrangement is not possible in the Pfr tertiary structure here reported because of steric clashes between the PAS9 domains in the contracted position and the changes in the helical spine trajectory (GAF-PHY and PHY-OM helical linkers). Nevertheless, to evaluate the feasibility of the crystallographic tongue-settled antiparallel assembly, adaptive Poisson-Boltzmann solver (APBS) electrostatic calculations were performed and compared to a projected parallel head-to-head model (see details in Materials and Methods section). The results show that the configurational and binding energies are similar in both quaternary arrangements and within the range of a stable dimer (**Figure S17 and Table S3**).

In order to probe the existence of different dimeric arrangements in solution for the Pr and Pfr states, we performed SEC experiments with wild-type, G454E and G454A holoproteins under far-red light or dark conditions, maximizing the relative abundance of these photostates, respectively. All variants showed a single and symmetric peak and present almost identical elution fractions that vary with the light conditions. The three samples illuminated with far-red light (prior to and during the SEC run) showed a clear shift toward lower elution fractions compared to the dark-adapted samples (**Figure 6B**). Additionally, the peak of the control variant D199A, which does not photoconvert and is locked in a Pr-like state (*40*), overlaps with the peaks of the far-red irradiated bathy samples, regardless of the light treatment (**Figure 6B**). Altogether, these results are in agreement with *Xcc*BphP transitioning between two dimeric assemblies in response to light.

Finally, we designed a cross-linking strategy based on the differential disposition of the residues at the protomer-protomer interface between the two dimer configurations to demonstrate the existence of the Pfr antiparallel quaternary arrangement in solution (**Figure 6C**). We used bissulfosuccinimidyl suberate (BS3), a cross-linker that reacts with primary amines (*i.e.* side chains of Lys residues and N-termini) because, according to the crystallographic quaternary assembly of the full-length Pfr structure, two pairs of Lys are within the cross-linking distance range, namely Lys451/Lys460’ and Lys451’/Lys460 (**Figure 6C, *upper panel* and Figure S18**). Accordingly, the presence of a cross-linked dimer is indicative of the existence of the antiparallel Pfr dimer. Wild-type dark-adapted or far-red irradiated holoprotein samples were incubated in the presence of a BS3 molar excess, then the samples were resolved in a denaturing SDS-PAGE, breaking non-covalently linked dimers. A band corresponding to the cross-linked dimer was observed exclusively in the dark-adapted samples -enriched in the Pfr form-, whereas the samples irradiated with far-red light -enriched in the Pr form-only showed the band corresponding to monomers (**Figure 6C**). This experimental evidence is in full agreement with the presence of a head-to-tail arrangement described in the Pfr structure and the light-gated transition between the two dimeric assemblies.

## DISCUSSION

Despite the vast structural information reported on photoreceptors, their dynamic protein structures pose a challenging matter in structural photobiology. To date, some valuable studies have reported structural data of a full-length phytochrome in both its Pr and Pfr photostates. From those, the Vierstra’s and Westenhoff’s groups have revealed models (at >10 Å resolution) of the two forms of *Dr*BphP by EM (*19*, *41*) and SAXS (*42*), respectively, but with different proposed mechanisms for the OM activation (“opening model” and “rotational model”, respectively). On the other hand, the Winkler’s group has recently solved the crystal structures of the *Idiomarina sp. A28L Is*PadC in both photostates, from which another mechanism for the OM activation has been proposed (“register model”); however, although the Pr state was crystallized as a homodimer (*43*), the Pfr state was defined in a heterodimeric assembly (one promoter in Pfr and the other one in Pr) derived from a constitutively active mutant variant (*35*).

The lack of full-length structures solved at the (near-)atomic resolution in both pure (non-mixed) Pr and Pfr states leaves gaps in the structural mechanisms involved in the signal transmission pathways during phytochrome photoconversion. The *Xcc*BphP crystal structures reported here in both photostates, combined with experimental and computational studies, allowed us to describe a more complete and precise landscape on the light-driven conformational changes from the chromophore to the OM during the reversible photoswitching in a full-length phytochrome.

The structural changes in the chromophore-binding pocket and the tongue region upon the BV isomerization from the Pr *ZZZssa* to the Pfr *ZZEssa* configuration agreed well with those reported in other BphP structures solved in each photostate (*8*, *9*). From those, definitely the most notable features are the flip-and-rotate movements of two highly conserved tyrosine residues surrounding the chromophore ring D (Tyr168 and Tyr195 in *Xcc*BphP) along with the tongue fold adopting an antiparallel β-sheet conformation in the Pr state and an α-helical structure in the Pfr state. This tongue fold interconversion, elegantly confirmed in the same BphP using dark and illuminated crystal structures of the PSM from *Dr*BphP (*22*), is a hallmark contrasting both structural photostates.

The extended protein elements associating the photo-induced changes in the chromophore-binding pocket with the tongue conformation have not been fully characterized yet. Despite this, a series of structural rearrangements that associate both regions in the different photostates have been exposed in most of the BphP structures solved to date (*8*, *9*). Our structures are in complete concordance with these structural features, confirming that the contrasts between the Pr and Pfr structures previously reported are preserved in a full-length BphP. Among these, the most relevant ones involve the aspartate residue from the DIP motif (Asp199 in *Xcc*BphP) located in the chromophore-binding pocket, which is found to be interacting with the arginine and the serine residue from the highly conserved PRxSF motif at the PHY tongue (Arg472 and Ser474 in *Xcc*BphP), in the Pr and Pfr states, respectively. This interdomain contact swapping is accompanied by an interchange in the position of two conserved tryptophan residues from the motifs W-G/A-G (Trp452 in *Xcc*BphP), and WxQ (Trp478 in *Xcc*BphP) from the Trp switch (*38*).

According to our results, the residues Leu193 and His194, which dwell outside the chromophore-binding pocket but close to Tyr195 (from the tyrosines flip-and-rotate mechanism), are also associative structural elements between these regions. The Leu193 side chain movement, driven by the flipping of Tyr195, seems to affect the hydrophobic pocket settled in the tongue-GAF interface observed in the Pfr state. We postulate that this structural rearrangement, not previously described for other BphPs, is interconnected with the hydrophobic area fashioned around ring D by the conserved tyrosine pair. Interestingly, *Xcc*BphP polar variants at position 193 (*i.e.* L193H, L193T, L193R, L193E, and L193C) produced Pr-favored proteins, while hydrophobic substitutions (*i.e.* L193F and L193M) generated Pfr-favored variants (*34*). This supports the observation that the substitution of this residue by polar ones may cause a destabilization of the inter-domain hydrophobic pocket, arranged in the Pfr state. Remarkably, leucine is the most frequent residue present at the homologous position from Leu193 in other phytochromes (**Figure S19**), indicating that the above mechanism might be shared among them. This position has been previously described to be key in the Pr-to-Pfr photoconversion by other groups. Yang *et al.* have shown in the bathy-type *Pa*BphP, that a substitution in the homologous position (Gln188) by a leucine residue affected its photocycle producing a mixed Pr-Pfr state (*12*, *18*). In the same line, Burgie *et al.* have confirmed that substitutions of the homologous residue in *Dr*BphP (His201) dramatically compromises its full Pr-to-Pfr photoconversion (*24*). More recently, a study on the PSM from Agp2-PAiRFP2, a fluorescence optimized BphP derived from *A. fabrum* Agp2, has revealed that Phe192 flipping (Tyr195 in *Xcc*BphP) upon BV isomerization produces a positional shift of Gln190 (Leu193 in *Xcc*BphP), which would induce a tongue refolding to the Pr state by steric hindrance with Trp440 (Leu458 in *Xcc*BphP) (*28*). Interestingly, this mechanism might share some analogous features to the one proposed here for *Xcc*BphP, since Leu458 is structured inside in the hydrophobic pocket in the Pfr state and disordered in the Pr state (**Figure 4B**).

On the other hand, during the Trp switch, His194 is found to be interacting with Trp452 in Pr or Trp478 in Pfr. This interaction seems to be more crucial in the Pfr state as revealed by the photochemical behaviour of the H194A variant (**Figure 1 and Table 1**). As His194 can act as a cation or as a π-system depending on its protonation state, the nature of its interaction cannot be assigned solely based on the structural data. Nevertheless, positive residues or residues carrying a partial positive charge are mostly found at the homologous position of His194 in phytochromes (**Figure S19**). Thus, it is reasonable to propose that the contact between this position and the aromatic residues from the tongue motifs W(G/A)G and (W/Y/F)x(E/Q) may be a conserved cation-π interaction. However, further research is needed to evaluate the nature of this GAF–tongue contact and its influence on downstream signal transduction.

Altogether, our structures suggest that residues −2 and −1 from the second (from N-to-C terminus) flipping tyrosine position may be important linkers between the tongue and the chromophore-binding pocket during the Pr-to-Pfr photoconversion.

Undoubtedly, one of the most relevant concerns is how the long-range structural changes are propagated from the PSM to the OM during the Pr-Pfr photoconversion. There is a broad consensus that the tongue fold interconversion and the associated structural motions affect the PHY domain position by pull/push movements, and thus the main helical spine trajectory (*9*, *22*). This large-scale structural rearrangement in the PSM, also termed “toggle model” (*7*, *29*), is proposed to be essential for modulating the OM position and its activation/deactivation after the photoactivation cascade. However, this notion mostly arises from structures derived from truncated BphP versions (without a complete OM). A recent characterization of the PSM from *Dr*BphP by solution NMR experiments proposed a pathway for the signal transduction from the chromophore binding-pocket to the helical spine *via* the figure-of-eight knot along with the PAS and GAF domains (*44*). Interestingly, these regions remain invariable in the Pr and Pfr structures here reported. Thus, this mechanism is absent or is consigned to intermediate states during the *Xcc*BphP photoconversion.

Our structures reveal a prominent reorientation in the PHY position with alterations in the GAF-PHY and PHY-OM helical linkers along with a likely inversion of the quaternary assembly when contrasting both states. Indeed, helical deviations have been frequently reported in sensory histidine kinases, comprising photoreceptors, which are crucial for protein signaling (*45*, *46*) and helical hinges are common in transmembrane proteins, being associated with their functional dynamics (*47*).

The GAF-PHY helical linker has been proposed to be correlated with the photostate, being straight in Pfr and curved in Pr (*9*, *48*). We observed the same structural behavior in both full-length photostates, where the Pr structure shows a helix bending (kink 1). Remarkably, the comparison between the Pfr structures from G454E and ΔPAS9(1-511) also display a torsion (albeit less pronounced), which indicates that this region has a particular flexibility and may be affected by the OM omission. Thus, changes in the GAF-PHY helical linker from PSM fragments should be cautiously analyzed when comparing different photostates.

The PHY-OM helical linker connects the PSM with the OM, and thus it is a key structural element for coupling the light perception and the allosteric response. In this region, we noted three variations between both photostates, a change in the trajectory (hinge), a helical bending in Pr (kink 2), and an unprecedented helical disruption in Pfr (break). The analysis of the *Xcc*BphP structures revealed that the PHY-OM helical linker trajectory is indeed altered by the push/pull effect on the PHY domain generated by the tongue interconversion, validating the toggle model in a full-length BphP (please refer to **Figure 4D** for the structures aligned at the GAF domains, and to **Figure 5A**, right panel, for the structures aligned at the PHY domains).

As the Pr dimer interface is largely arranged on the helical spine, these helical rearrangements on the GAF-PHY and PHY-OM helical linkers should partially or completely destabilize the head-to-head dimeric assembly and, consequently, open it. In fact, strong and crucial Pr dimer interactions such as Glu328/Arg501 and Glu505/Arg509 stabilizing the GAF-PHY/PHY-OM and PHY-OM/PHY-OM helical linkers, respectively, are switched by intramolecular interactions in the Pfr state. This scenario is also supported by the Pfr structures of the PSM from *Dr*BphP and the (near) full-length *Rp*BphP1, which showed an open dimer conformation (*22*, *33*). Moreover, previous works on *Dr*BphP revealed that some of the dimers were opened at the histidine kinase region (*19*) and that the dimerization interface between the OMs could be broken by light absorption (*49*). Therefore, the dissociation of the dimer interface is a possible way of action during photoconversion. In this regard, the contribution of the OMs to the dimer interface in the Pr head-to-head quaternary assembly would respond to why ΔPAS9(1-511) mainly behaves as Pfr. Additionally, the two point subtitutions in the R525E-R527E variant, which were designed to disturb the Pr dimer interface, generated a monomeric full-length Pfr-favored protein, showing that the interactions between the PHY-OM helical linker and the PAS9 domain from neighboring protomers are crucial for the Pr dimer stabilization. Accordingly, our computational analysis showed lower stability on the PHY-OM helical linker in a hypothetical monomeric scenario, compared to the dimer, which would facilitate the unprecedented disruption (break) in the helical spine, and subsequently generate an abrupt change in the OM position.

The breaking point of the PHY-OM helical linker is located at Phe512. Interestingly, this residue has been previously identified by our group as a potential key structural element in the light-driven conformational changes of *Xcc*BphP (*31*). Aromatic residues have high propensity to be on helix kinks, and interestingly, the (F/Y/W)xxx(F/Y/W) motif (_512_FQQDF_516_ in *Xcc*BphP) has been frequently observed in kinked or broken helices (*50*). Moreover, the probability of helical disruption for phenylalanine is larger than for other aromatic residues (*51*). It is unclear how the helical breaking is established. A possible scenario is that the abolition of the Arg436-Phe512 cation-π interaction, triggered by the PHY domain relocation in the Pfr state, provokes an exposition of the Phe512 hydrophobic side chain to the solvent. This change in the hydration environment may generate a conformational rearrangement in Phe512 that breaks the hydrogen bond between its carbonyl group and the Phe516 (*i* + 4) amide group of the helical turn, and thus introducing a destabilizing kink in the α-helix.

The combined structural motions described here might be followed by a dimer arrangement switch from head-to-head settled on the helical spine in Pr to head-to-tail settled on the tongue in Pfr. The functional dimerization assembly in this kind of photoreceptors is still controversial, but could be crucial in the regulation of the OM activation. Although parallel assemblies are the most observed in BphPs, antiparallel arrangements have also been reported (*11*, *28*, *33*, *38*, *48*, *52*, *53*). However, in both cases, the dimer interface is typically arranged on the helical spine from both protomers. The parallel/antiparallel quaternary remodeling between the Pr and Pfr states here reported and supported by spectroscopic, biophysical, biochemical and computational approaches, associates three major structural players of BphP upon photoconversion: the tongue, the helical spine, and the OMs. Additionally, the remodeling is in line with a proposed signal output (de-)activation in which both OMs might be needed to work in concert or separated in the different photostates (*7*).

Recently, Takala and colleagues summarized the three probable structural mechanisms for OM activation (*9*): (i) opening model (*19*, *41*), where a partial opening of the dimeric arrangement might take place, (ii) rotation model (*42*), which may involve a rotation of the dimer around the helical spine, and (iii) register model (*35*, *43*), which might affect the helical spine register. According to our results, the *Xcc*BphP photoactivation may contain elements of the “opening model” in the aperture of the Pr dimer (“Y-shaped dimer”), followed by a new defined “joint-inversion model” to explain the events involved in the PHY-OM helical linker and the quaternary reassembly. As a whole, we propose a general mechanistic model for the photoconversion in *Xcc*BphP, starting in the photoreaction center and culminating with the long-range allosteric reorientation of the OM (**Figure 7**). The main series of events comprises (i) the ring D isomerization of the BV molecule upon the absorption of a photon or dark conversion, (ii) the reorganization of the networks within and around the chromophore-binding pocket, (iii) the β-sheet/α-helix tongue transition, (iv) the PHY reorientation, with the concomitant GAF-PHY and PHY-OM helical linkers changes, (v) the dimer dissociation, (vi) the PHY-OM helical linker rectify/breaking with the reorientation of the PAS9 domain, and (vii) the parallel/antiparallel dimer inversion assembly.

**Figure 7.**
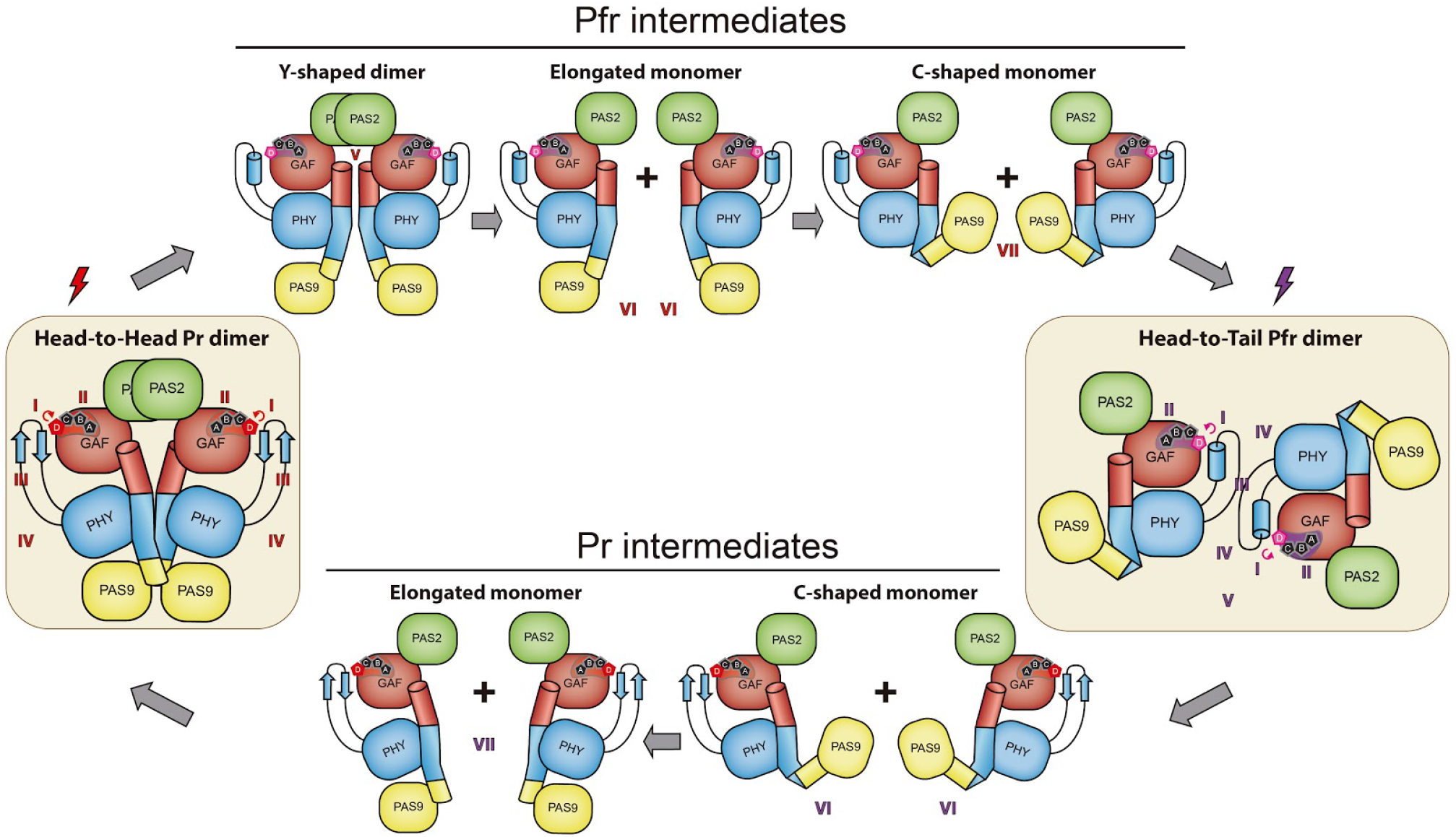
Proposed *Xcc*BphP photoconversion model. The Pr and Pfr crystal structures are schematically represented on the left and the right yellow panels. The hypothetical intermediates involved in the Pr-to-Pfr and the Pfr-to-Pr photoconversions are depicted. Domains are colored according to **Figure 2**. The red and the far-red irradiations are represented as red and purple lighting cartoons, respectively. This proposed mechanism starts in the photoreaction center and culminates with the long-range allosteric reorientation of the OM. The main series of events comprises (i) BV ring D isomerization upon photon absorption or dark conversion, (ii) reorganization of the networks within and around the chromophore-binding pocket, (iii) β-sheet/α-helix tongue transition, (iv) PHY reorientation, with the concomitant changes in the GAF-PHY and PHY-OM helical linkers, (v) dimer dissociation, (vi) PHY-OM helical linker rectification/break with the reorientation of the PAS9 domain, and (vii) parallel/antiparallel dimer inversion assembly. In the Pr-to-Pfr photoconversion, the BV ring D isomerizes to the *ZZEssa* configuration followed by an α-helix conformational transition of the tongue. As a result, a push movement of the PHY domain along with a straightening of the GAF-PHY and PHY-OM helical linkers would trigger the detachment of the intertwined OMs (PAS9) generating a Y-shaped intermediate. The loss of dimeric interactions dissociates the intermediate into monomers. Upon dissociation, the PHY-OM helical linker suffers a break, generating a C-shaped monomer. Then, these monomers find a new dimerization surface, through the tongue in the α-helix conformation into an antiparallel dimer. The Pfr-to-Pr photoconversion starts by ring D isomerization to the *ZZZssa* configuration, followed by a β-sheet transition of the tongue, disrupting its dimeric interface hindering dimer intermediates, and producing C-shaped monomers. As a result, the PHY domain is pulled along with the bending of the GAF-PHY and PHY-OM helical linkers. Next, PHY-OM helical linker suffers a rectification, reconstituting the helical spine dimeric interface, and ultimately the formation of the Pr dimer.

As the PSM overall architecture and the helical spine are conserved in several phytochromes, the mechanistic features exposed in this work can help to elucidate the downstream elements triggered by light in other members of this photoreceptor family, which are required to transduce the signal from photoreception to the biological response. However, as the nature of the OMs is variable, and consequently, their extended dimerization interfaces might have strong regulatory differences for the long-range signal transduction, diverse OM photoactivation mechanisms can occur depending on their specific function, and thus preventing a universal common principle.

## MATERIALS AND METHODS

### Generation and purification of recombinant *Xcc*BphP variants

The pET24a-*Xcc*BphP expression vector coding for the wild-type version of the *Xcc*BphP protein (ORF XC_4241) from *Xanthomonas campestris* pv. *campestris 8004* is from previous work (*54*). The cloned ORF includes an N-terminal methionine and His-tag followed by the complete coding sequence (with the exception of its starting residue: 2–634), totaling 640 residues. Some mutant constructs were generated using pET24a-6xHis-*Xcc*BphP (*32*) as a template, through a Gibson assembly approach (namely D177A, H194A, S280V, W452A, W452A-G454E, A453E, W478A, R501A, F512A and F512P). Briefly, two independent amplicons, with 3’ and 5’ complementary regions, were generated from the template using specific primers (**Table S1**, one set of primers bearing the specific mutation, while the other corresponds to the kanamycin cassette on the vector backbone). Then, both PCR fragments were assembled following the Gibson assembly protocol, where only whole reconstituted vectors were able to transform bacteria. The R525E-R527E mutant plasmid was generated using pET-24a-XccBphP as a template for mutagenesis using whole plasmid amplification followed by DpnI template digestion (New England Biolabs). This was achieved using Q5 High-Fidelity DNA Polymerase (New England Biolabs), the corresponding set of specific primers (**Table S1**) and digestion of the template from the reaction mix with DpnI (New England Biolabs), following the manufacturer’s instructions. The mix was later used to transform bacteria. Finally, the vectors coding for the truncated versions ΔPAS9(1-525) and ΔPAS9(1-527) were generated first, by amplifying the pET-24a-*Xcc*BphP template by PCR using specific primers (**Table S1**). The resulting amplicons were digested with NdeI and BamHI and ligated with T4 Ligase (New England Biolabs) into a pET-24a empty vector digested with the same restriction enzymes. The ligation was used for bacterial transformation. All cloning was performed in *Escherichia coli DH10B*, the resulting mutations were confirmed by Sanger’s DNA sequencing. The constructs corresponding to pET-24a-*Xcc*BphP D199A, V266S, G454E, G454A, S474E and ΔPAS9(1-511) were obtained in previous works (*31*, *34*). All resulting vectors were used to transform *E. coli BL21(DE3)pLysE* for recombinant protein production.

Bacterial strains harboring each expression vector were cultured at 37 °C in Luria–Bertani medium. When required, the antibiotics kanamycin (Km) and chloramphenicol (Cm) were added in final concentrations of 35 and 25 μg mL^−1^, respectively, and induced with a final concentration of 0.5 mM IPTG overnight at 20 °C with agitation (250 r.p.m.). Cells were harvested, ruptured by sonication, and apoproteins and holoproteins were purified by means of nickel–NTA affinity and size-exclusion chromatography steps as described previously (*32*). For the case of holoproteins a 1-h incubation with BV at room temperature in the dark (Sigma-Aldrich) was performed before size-exclusion chromatography. Protein concentration was estimated using the calculated molar extinction coefficients at 280 nm provided by the ExPASy ProtParam tool (https://web.expasy.org/protparam/) based on the polypeptide sequences. In the UV-Vis spectroscopy experiments, the protein buffer composition was 50 mM Tris-HCl, 250 mM NaCl, pH 7.5. For crystallization, the final protein buffer composition was 10 mM Tris-HCl, 25 mM NaCl, pH 7.6.

### UV-Vis spectroscopy and data analyses

Dark-assembly of wild-type *Xcc*BphP and G454E was performed by incubating the apoproteins with BV in a 2:1 molar ratio inside quartz cuvettes, in 50 mM Tris–HCl, 250 mM NaCl, pH 7.5 at 25 ºC. UV-Vis spectra were recorded in the dark since the addition of BV and the spectra were collected every 12 s for 0-15 min, every 30 s for 15-30 min, every 1 min for 30-45 min, every 2.5 min for 45-60 min, and every 10 min for 60-180 min. All UV-Vis measurements in this work were performed in a spectrophotometer (Model Cary 60, Agilent Technologies) with an installed temperature control Peltier module (**Figure S1**).

Dark conversion experiments were initiated in quartz cuvettes containing holoprotein solutions of *Xcc*BphP or its variants at ~1 mg mL^−1^ in 50 mM Tris–HCl, 250 mM NaCl, pH 7.5 at 25 ºC. Samples were irradiated for 20 min with red LED light (630 nm, fluence 0.2 μmol m^−2^ s^−1^) or for 7 min with far-red LED light (733 nm, fluence 0.6 μmol m^−2^ s^−1^). Sequential absorption spectra were recorded in the dark (**Figure S2**).

The pure-Pr and pure-Pfr spectra were calculated using data derived from the dark-conversion datasets initially illuminated with far-red light and/or red light. The spectra most enriched in Pr (highest Abs_684nm_:Abs_754nm_ ratio) or Pfr (lowest Abs_684nm_:Abs_754nm_ ratio) in the variant datasets were selected (Pr-enriched and Pfr-enriched, respectively). The pure-Pfr was calculated similarly as previously described by Assafa *et al.* (*55*). Briefly, a series of spectra were generated by subtracting the Pfr-enriched spectrum with increment fractions (0 to 1 with increment steps of 0.001) of the Pr-enriched spectrum. The derivative spectra of the subtraction series were subsequently calculated, and the minima and maxima around the wavelength corresponding to the Pfr maximum were determined. Finally, the pure-Pfr was estimated as the subtraction that minimized the difference of the derivatives corresponding to the above-mentioned minima and the maxima wavelengths (**Figure S3A**).

The pure-Pr was estimated, first, by calculating a series of spectra subtracting the Pr-enriched spectrum with increment fractions (0 to 1 with increment steps of 0.001) of the pure-Pfr spectrum. Then, the derivative spectra of the subtractions were subsequently calculated, and for each one of them a mean and standard deviation (SD) were calculated in the range between the wavelength corresponding to the Pfr maxima and the highest wavelength in the dataset (820 nm). Finally, the pure-Pr spectrum was calculated by the subtraction of the pure-Pfr fraction corresponding to the local maxima produced by the multiplication of the mean and SD values of the derivative spectra in the selected range (**Figure S3B**).

Then, for each spectrum, assuming to be composed of two pure components, we calculated the linear combinations of their pure-Pfr and pure-Pfr spectra to best fit the dataset using data corresponding to wavelengths above 510 nm. This allowed us to obtain the individual Pr and Pfr spectra at each time point that added together reconstituted the experimental dataset (**Figure S4**). The Pr and Pfr abundances (%) at each time were estimated as the relative spectral area corresponding to the pure-Pr and the pure-Pfr with respect to the sum of the areas (using the data corresponding to wavelengths above 510 nm).

The kinetic analysis was carried out by optimizing the parameters of a double exponential model (Eq. 1) and a mono exponential model (Eq. 2) to fit to the Pr / Pfr abundance data. The Pr or Pfr equilibrium values were derived from the parameter A from the double exponential fitting, while the mono exponential model was used to estimate half-life values using the parameter H (**Table 1**).

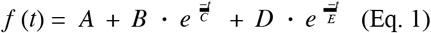

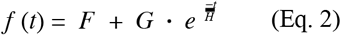

As an example, a processed dataset for the wild-type version is available in GIF format as **Supplementary Information**, overlaying individual pure-Pr and Pfr spectral components, and their relative abundances at each time point. All steps of the analysis were performed using custom made scripts developed in MATLAB software (version 2015b).

### SLS-SEC measurements

The average MWs of *Xcc*BphP and its variants were determined using a PD2010 90° light scattering instrument (Precision Detectors) connected in tandem to a high-performance liquid chromatography system (HPLC) and a LKB 2142 Differential Refractometer Detector (Pharmacia). Only MWs for apoproteins could be calculated due to the interference produced by BV in the 90° light scattering measurements because of its absorption in the laser wavelength (685 nm, AlGaInP). A Superdex 200 HR-10/30 column (24 mL; GE Healthcare Life Sciences) was used with isocratic elution in 50 mM Tris-HCl, 250 mM NaCl, pH 7.7 at a flow rate of 0.4 mL · min^−1^ at 20 °C, with 0.08-0.22 mg of injected protein sample. The MW was calculated relating its 90° to the IR signals using the software Discovery32 supplied by Precision Detectors. Bovine serum albumin (BSA, MW 66.5 kDa) was used as a standard.

### SEC elution fraction determination

Holoprotein samples of wild-type *Xcc*BphP and its variants G545E, G454A and D199A were incubated for 24 h in the dark, and then either kept in the dark or irradiated for 7 min with a far-red LED source (733 nm, fluence 0.6 μmol m^−2^ s^−1^) prior to the experiment. The dark-adapted samples were subjected to FPLC-SEC at room temperature in a Superdex 200 HR-10/30 column. The total amount of the protein samples injected in the column was 0.1 mg. The column was covered in aluminium foil for the dark experiments, while the irradiated samples were kept under far-red light during the SEC. The SEC buffer was 50 mM Tris-HCl, 250 mM NaCl, pH 7.5 at a flow rate of 0.6 mL · min^−1^ and the recorded signal was absorbance at 280 nm. Data were normalized and plotted using GraphPad PRISM software. The manipulation of the dark samples was performed under dim green-filtered light using a green acetate band-pass filter (LEE lighting, cat. 090) with a transmission maximum at 525 nm.

### Dimer Cross-Linking and electrophoresis

Dark-adapted or far-red irradiated (pre-irradiation for 7 min and irradiation during the experiment with a far-red LED source of 733 nm, fluence 0.6 μmol m^−2^ s^−1^) *Xcc*BphP wild-type samples at 1 mg mL^−1^ (14 μM) were incubated with bissulfosuccinimidyl suberate (BS3) (Thermo Fisher Scientific) in 250, 125, 100 and 25 molar excess for 100 for 30, 60 and 120 min at 22 °C in a buffer containing 50 mM phosphate, 250 mM NaCl, pH 7.5. All cross-linking reactions were quenched with 100 mM Tris-HCl, pH 7.5 (final concentration). Then, the loading buffer was added to the samples and resolved by SDS-PAGE (10% gel). The manipulation of the dark samples were performed under dim green-filtered light using a green acetate band-pass filter (LEE lighting, cat. 090) with a measured maximum at 525 nm. The BS3 crosslinking distance range is 11 Å according to the manufacturer.

### Crystallization

*Xcc*BphP was crystallized as reported previously (*32*). However, a slower adaptation of the crystals to the cryoprotectant before the flash-cooling into liquid nitrogen was crucial to reach a better resolution of the diffraction data in comparison with the existing structure (PDB entry 5AKP). For the ΔPAS9(1-511) construct and the G454E mutant, initial crystallization conditions were screened on sitting-drop Greiner 609120 96-well plates using a Honeybee963 robot (Digilab) and commercial kits from Jena Bioscience (Jena) and Hampton Research (Aliso Viejo) at 20 and 13 mg · mL^−1^, respectively. In both cases, thin green bars appeared after a few weeks of equilibration at 21 ºC in several solutions of the kits. Some of these initial conditions were then optimized in 24-well hanging-drop Hampton Research VDX plates. In this sense, diffraction-quality ΔPAS9(1-511) crystals were grown with a precipitation solution consisting of 6% (*w*/*v*) polyethylene glycol (PEG) 4000, 0.1 M HEPES, 10% (*v*/*v*) isopropanol, pH 6.7, whereas G454E crystals appeared with 30% (*w*/*v*) pentaerythritol propoxylate (5/4 PO/OH), 0.1 M MES, pH 6.8. Samples were cryoprotected in their respective mother liquors added with 29% (*w*/*v*) PEG 400 for ΔPAS9(1-511) and 5% (*w*/*v*) PEG 400 for G454E, and then flash-cooled in liquid nitrogen using synthetic mounted cryoloops (Hampton Research). In all cases, crystallization trials were performed in the dark, while crystal handling and vitrifying procedures were carried out under dim green-filtered light using a green acetate band-pass filter (LEE lighting, cat. 090) with a measured maximum at 525 nm. Crystals were observed under the microscope using a 3-W power 520 nm green LED light filtered with the same green acetate band-pass filter.

### Data collection, structure resolution and refinement

X-ray diffraction datasets were collected at the PROXIMA-1 and PROXIMA-2A beamlines at synchrotron SOLEIL (France) on several crystals with the help of the MXCuBE application (*56*) using the classical and helical modes. Datasets were indexed, integrated and scaled with XDS (*57*) and AIMLESS (*58*), leaving 5% of the reflections apart for cross validation purposes. All structures were solved by the molecular replacement method with PHASER (*59*) using the coordinates of the wild-type protein as a search model (PDB entry 5AKP) (*31*), and checked for proper packing. Refinement and manual model building were then performed with BUSTER (*60*) and COOT (*61*), respectively. The 2*mF*_o_-*DF*_c_ electron density maps allowed for a fairly complete trace of the protein backbones with the exception for the initial 8 to 10 N-terminal residues, and the regions comprising the residues 457-469 for the wild-type protein, 393-398, 527-528, 601-606, 616-623 for G454E, and 333-340, 389-401 for ΔPAS9(1-511), which correspond to exposed loops. The final models were validated with MolProbity (*62*). **Table S2** shows the most relevant statistics on the data collection and processing steps, as well as the PDB deposition information.

### Molecular Dynamics (MD) simulations and electrostatic energy calculations

For the classical all-atom MD simulations, starting structures (wild-type Pr state, PDB entry 6PL0, and G454E Pfr state, PDB entry 7L59) were completed for missing loops (124-126 and 457-469, for 6PL0; 384-391, 519-520, 593-598 and 608-615 for 7L59) with the *ab initio* loop modelling tool of MODELLER (*63*). Both structures were initially simulated in monomeric states. All structures were solvated with a 10 Å octaedric TIP3P water box. Standard protonation states were assigned to titratable residues (Asp and Glu negatively charged, Lys and Arg positively charged). Histidine protonation was assigned to favour hydrogen bonds formation in the crystal structure. The protein system was modelled using Amber ff14SB as a forcefield (*64*), while for the parametrization of the BV chromophore, the electrostatic potential for the ligand was computed using Hartree-Fock and 6-31G* basis set as implemented in Gaussian 09 (*65*). Antechamber (*66*) with a RESP scheme was used to compute partial charges for the molecule. All systems were subject to a mild minimization protocol of 1000 steps to remove bad contacts between atoms. The systems were slowly heated up from 10 to 300 K in 100 ps with a linear ramp in an NVT ensemble. Afterwards, density equilibration was performed for 1 ns in an NPT ensemble. Simulations were run with Langevin thermostat with a collision frequency of 5 ps^−1^ while the pressure was kept constant with the Monte Carlo Barostat. During both the heating and the equilibration process a 1 kcal/mol•Å^2^ harmonic restraint on all C_α_ atoms was placed. All simulations were done with Particle Mesh Ewald to treat long-range electrostatic interactions with a cutoff of 9 Å. All molecular dynamic runs were done with a 2 fs time step employing the SHAKE algorithm to keep heavy atom-Hydrogen bonds at equilibrium distances. The GPU-accelerated version of pmemd (v18) was used to run the simulations (*67*). For each system, four independent 500 ns simulations were done. To analyze these MD runs, dihedral angle computations and clustering were performed with CPPTRAJ (*68*) while statistical analysis and plotting were done with R and ggplot2. For clustering, the dbscan algorithm was used with an epsilon of 0.9 Å.

To assess dimer stability of the Pfr state in the two conformations, head-to-tail and head-to-head, we performed 500 ns of Gaussian accelerated MD (GaMD) (*69*) simulation. While the initial head-to-tail dimer model was constructed using the G454E full-length crystallographic structure, the head-to-head model was designed *via* homology modeling taking as a template the full-length dimeric assembly in the Pr state. Here, the PSM tertiary fold corresponds to that reported for the G454E monomer, and the PAS9 was rearranged in the same conformation as found in the Pr state, thereby obtaining a full-length Pfr head-to-head dimer similar to the Pr assembly. Simulations of *Xcc*BphP in aqueous solution were performed with Amber ff14SB, including BV parameters, as derived in a previous work (*70*). Minimization, heating and thermal equilibration were performed following the same protocols are described above for the *Xcc*BphP monomer models. In order to enhance conformational sampling, conventional MD simulations were extended by 500 ns of Gaussian accelerated MD (GaMD) simulations, with acceleration threshold σ_0_ = 5 (*69*), without any constraints. Electrostatic Energy Calculations on the two dimers were done with APBS (*71*), with 545 grid points in the x,y,z dimensions and a resolution of 0.3 Å on 500 snapshot structures derived from a 500 ns GaMD simulation.

### GAF sequence analysis

To generate the GAF domain sequence logo (**Figure S19**), the Seq2Logo web server (http://www.cbs.dtu.dk/biotools/Seq2Logo/) was used with default parameters (Kullback-Leibler logo, 0.63 for Hobohm1 clustering, and 200 for weight on prior / pseudo counts) (*72*). The input was a multiple sequence alignment (MSA) of a diverse and non-redundant dataset of GAF domain sequences from phytochromes, derived from previous work (*34*). Briefly, an initial phytochrome dataset was constructed with all sequences extracted from the UniProtKB and mapped UniRef50 database (for avoiding redundancy) and including the *Xcc*BphP sequence. Only sequences bearing a PAS2-GAF-PHY domain triad were selected. Subsequently, their corresponding GAF sequences were identified locally using HAMMER 3.2 software (http://hmmer.org/) and extracted using custom MATLAB scripts. This last dataset of 751 entries was used to construct the MSA using the Clustal-O program in the EBI server (https://www.ebi.ac.uk/Tools/msa/clustalo/) with default parameters. Additionally, absolute and relative frequency counts were performed on the MSA for positions corresponding to Leu193 and His194 of *Xcc*BphP.

### Structure representation

Molecular structures and their electron densities were represented using PyMOL Molecular Graphics System 1.8 (Schrödinger, USA). The electron density maps were generated in PHENIX (*73*).

## Accession Numbers

Coordinates and structure factors have been deposited in the Protein Data Bank (http://wwpdb.org/) with accession numbers 6PL0 (wild-type *Xcc*BphP, Pr state), 7L59 (G454E, Pfr state), and 7L5A (ΔPAS9(1-511), Pfr state).

## ACKNOWLEDGMENTS

This work was supported by the Argentinian Ministry of Science (MINCyT), the Argentinian Research Council (CONICET), and the National Agency for the Promotion of Science and Technology of Argentina (ANPCyT), under grants PICT 2015-0621 and PICT 2016-1425. LHO, SK, FAG, JR and HRB are researchers from CONICET. MSL is supported by an ANPCyT fellowship. LHO and SK acknowledge MINCyT for travel support. LAD is supported by the EMBL Interdisciplinary Postdoc Programme under Marie Curie Actions COFUND 664726. We are grateful for the access to the PROXIMA-1 and PROXIMA-2A beamlines at Synchrotron SOLEIL, France (proposal Nos. 20140791 and 20160817). We thank Dr. Francisco Velazquez-Escobar and Dr. Diego Ferreiro for their helpful advice.

## AUTHOR CONTRIBUTIONS

LHO, JR and HRB designed and supervised the project. LHO, MSL, SF, JR and HRB designed and performed the mutant constructs. SF, GTA, JR and HRB purified the proteins and performed UV–Vis spectroscopy experiments. HRB performed data analysis of UV-Vis data and bioinformatics. GTA, JR and HRB performed SEC–SLS experiments. JR analyzed SLS-SEC data. JR and SF performed the cross-linking assay. GTA, SS, JR and SK crystallized the proteins. LHO, SS, SK and LMGC performed X-ray diffraction data collection. LHO, SS and LMGC processed crystallographic data. LHO solved and refined the crystallographic structures. LAD, GB and MAM performed MD simulations. LHO, JR and HRB analyzed the protein structures. LHO, FAG, LMGC and HRB financed the project. LHO, JR and HRB wrote and illustrated the paper with contributions from all authors.

## Notes

The authors declare no conflicts of interest.

### Competing Interest Statement

The authors have declared no competing interest.

