## Supplementary Information for "Structural basis for the Pr-Pfr long-range signaling mechanism of a full-length bacterial phytochrome at the atomic level"

**The authors declare no conflicts of interest.**
Keywords: *Xanthomonas campestris*; photoreceptor; photosensor; phytochrome; bacteriophytochrome; BphP; biliverdin; red / far-red light; signal transduction; X-ray crystallography; UV-Vis spectroscopy; molecular dynamics

**Short title**
Pr-Pfr structures of *Xcc*BphP bacteriophytochrome

**Table of contents**

1. Figures S1-S19
2. Tables S1-S3
3. WT_Far-red.GIF
4. WT_Red.GIF

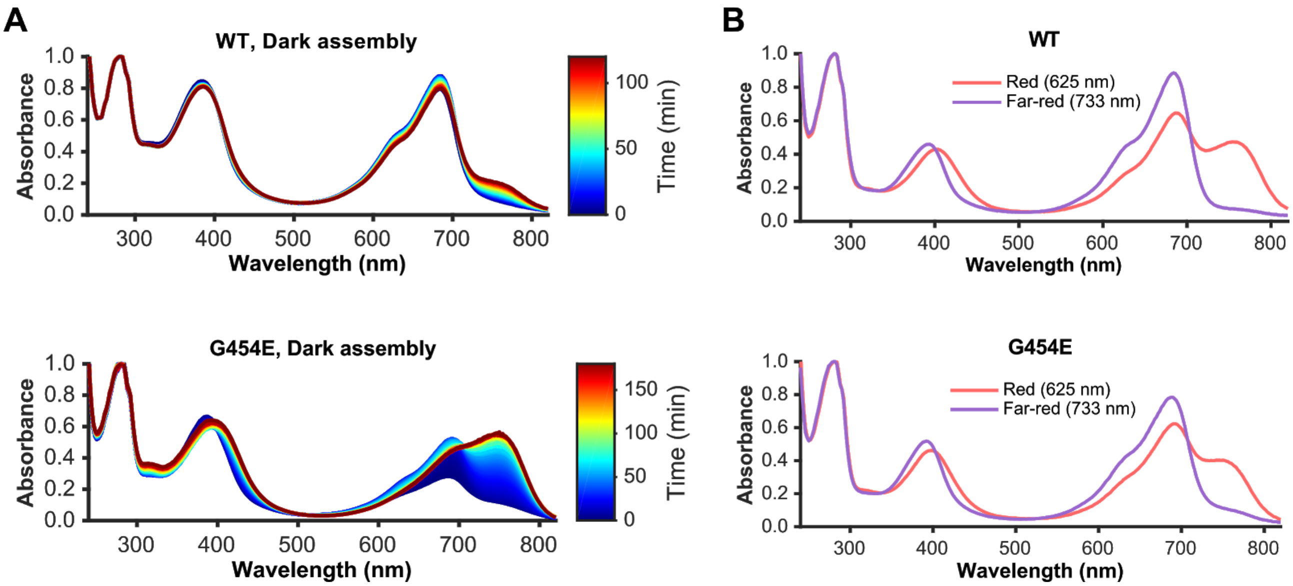


**Figure S1. Photochemical studies of *Xcc*BphP wild-type and G454E variants.**

A) Dark-assembly of wild-type *Xcc*BphP and G454E. *Xcc*BphP variants were incubated with BV in a 2:1 molar ratio. UV-Vis spectra were recorded since the addition of BV. B) UV-Vis spectra of wild-type *Xcc*BphP and G454E irradiated with red (625 nm) or far-red (733 nm) light. A-B) All spectra were normalized by the absorption values at 280 nm.

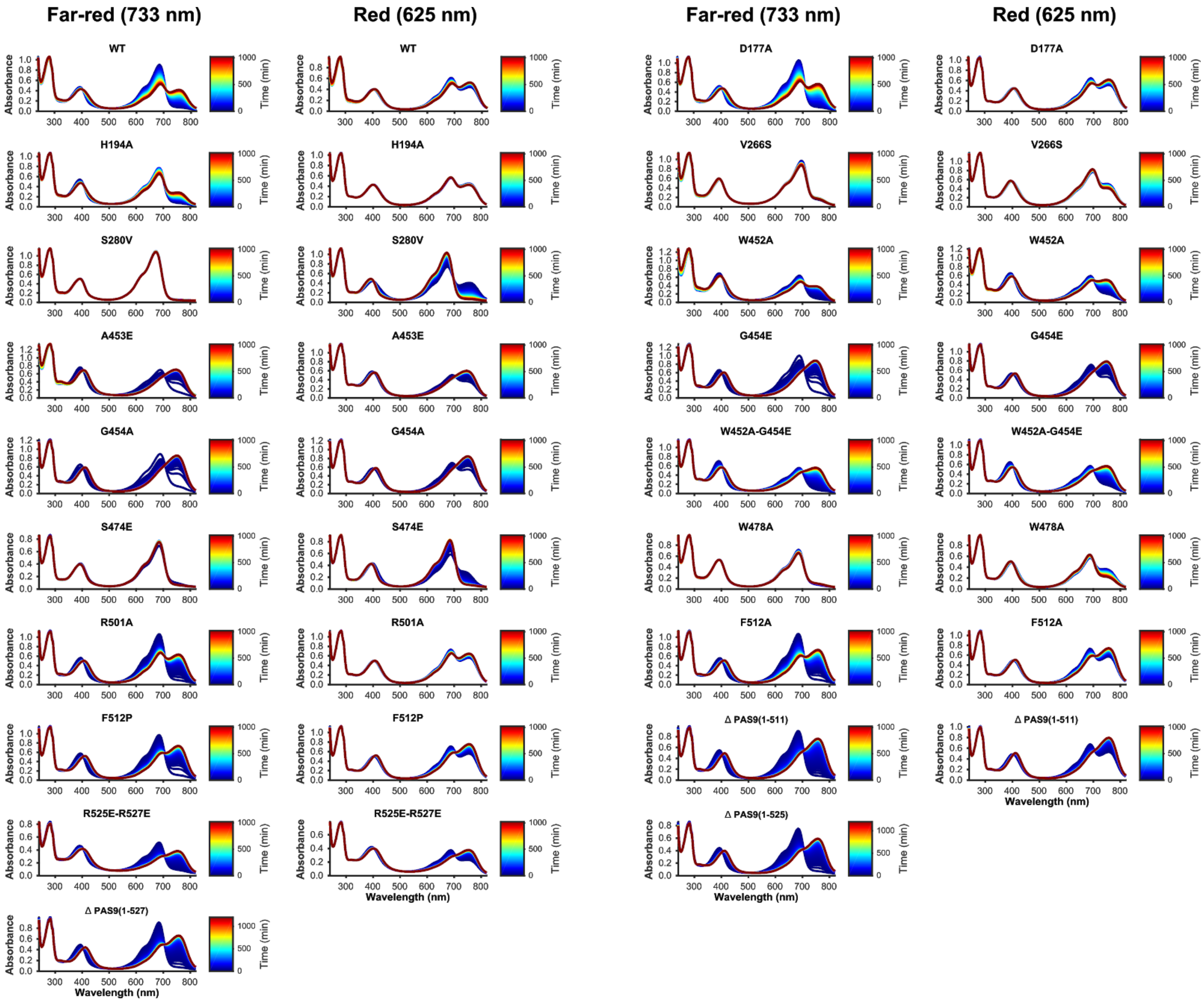


**Figure S2. Dark reversion experiments of *Xcc*BphP variants.**

Purified holoprotein samples (~1 mg $mL^{-1}$) were placed in a quartz cuvette, irradiated with red (625 nm) or far-red (733 nm) light, incubated in the dark and their UV-Vis spectra recorded over time (up to 960, 1020 or 1200 min). Spectra are represented in jet color code according to the time of acquisition.

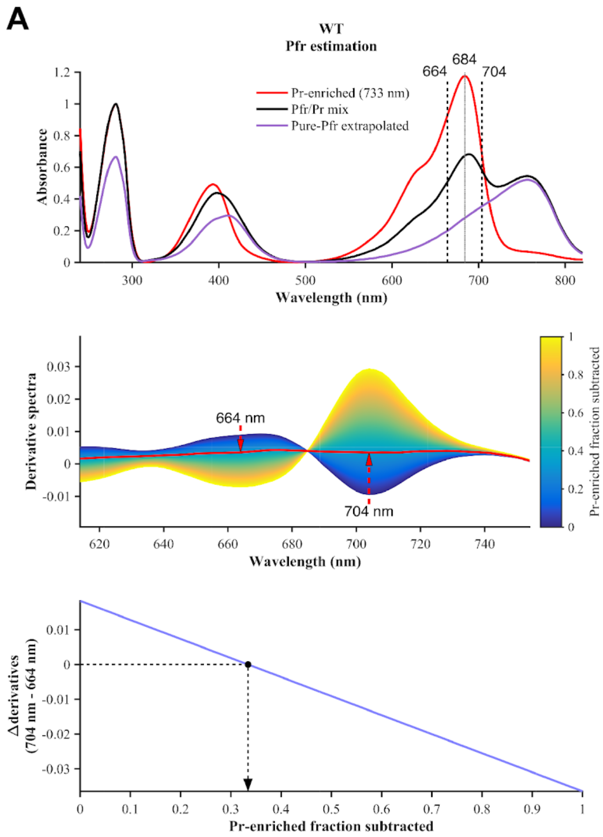


**Figure S3. Extrapolation of pure-Pr and pure-Pfr spectral components.**
The pure-Pr and pure-Pfr spectra were calculated using data derived from the dark-conversion datasets initially illuminated with far-red light and/or red light (top panels, Pr-enriched and Pfr/Pr mix spectra). A) Pure-Pfr estimation. A series of derivative spectra were generated after subtracting the Pfr-enriched spectrum with increment fractions of the Pr-enriched spectrum (middle panel). The minima and maxima around the wavelength corresponding to the Pfr maximum were determined (red arrows). The pure-Pfr was estimated as the subtraction that minimized the difference of the derivatives corresponding to the above-mentioned minima and the maxima wavelengths (bottom panel, the Pr-enriched fraction selected for the subtraction is indicated with a black dashed arrow). B) Pure-Pr estimation. A series of derivative spectra were generated after subtracting the Pr-enriched spectrum with increment fractions of the pure-Pfr spectrum (middle panel). The mean and standard deviation (SD) were calculated for each derivative spectra in the range between the wavelength corresponding to the Pfr maxima and the highest wavelength in the dataset (820 nm). The pure-Pr spectrum was calculated by the subtraction of the pure-Pfr fraction corresponding to the local maxima produced by the multiplication of the mean and SD values of the derivative spectra in the selected range (bottom panel, the pure-Pfr fraction selected for the subtraction is indicated with a black dashed arrow). Panels A and B show as example the pure-Pr and pure-Pfr extrapolations for the *Xcc*BphP wild-type. These procedures were performed to all variants in this study.

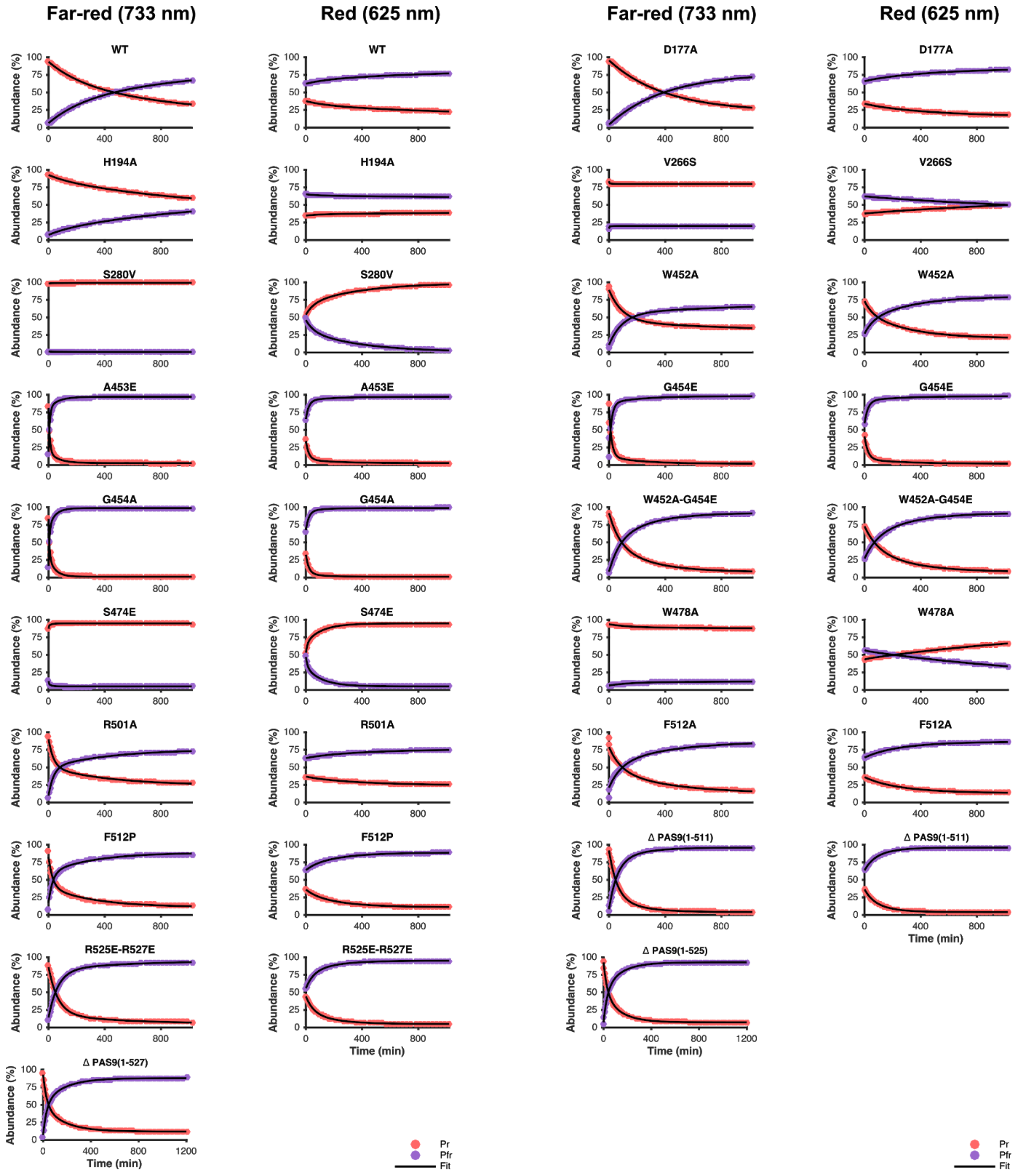


**Figure S4. Kinetic analysis of dark reversion experiments of *Xcc*BphP variants.**

The Pr and Pfr individual abundance (%) was calculated at each time for all variants using the processed data from **Figure S2** and the pure Pr and Pfr spectra estimated (**Figure S3**) and assuming that these two species are responsible for the absorbance changes during the experiment. Half-life values and Pr/Pfr abundance at equilibrium were estimated fitting equations 1 and 2 (black traces) to the resulting data, shown in **Table 1**.

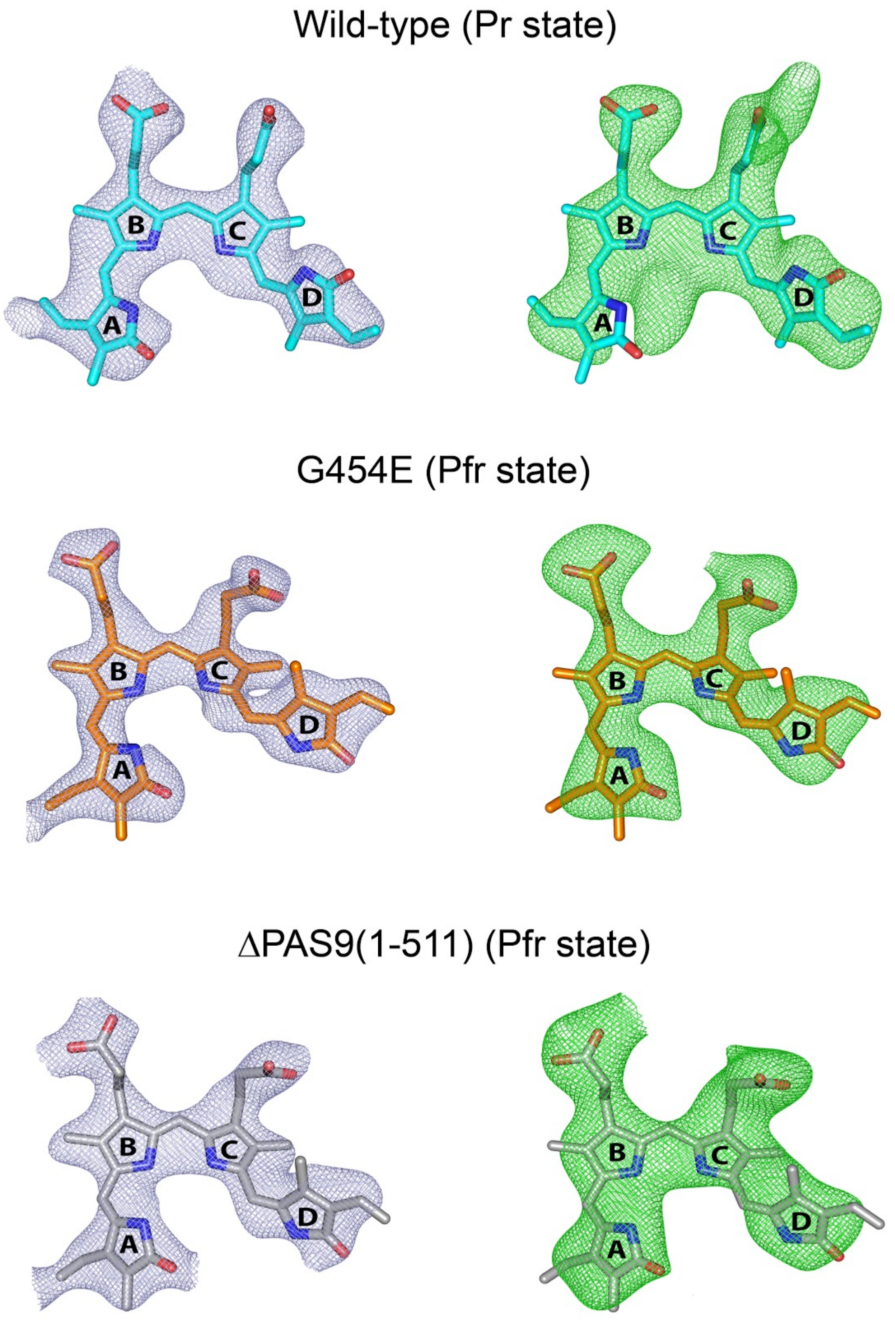


**Figure S5. Electron density maps around the BV chromophore at the *Xcc*BphP variants in the Pr and Pfr states.** Final $2mF_o–DF_c$ electron density maps (light blue mesh, left) contoured at the 1.0 σ level and final $mF_o−DF_c$ omit maps (green mesh, right) contoured at the 2.5 σ level. Omit maps were generated by removing the ligands from the models followed by five cycles of refinement in PHENIX (*73*). The BV molecule is shown as sticks with carbon atoms in cyan (wild-type), orange (G454E), and gray [ΔPAS9(1-511)], oxygen atoms in red, and nitrogen atoms in blue. The four BV pyrrolic rings are indicated as A, B, C and D.

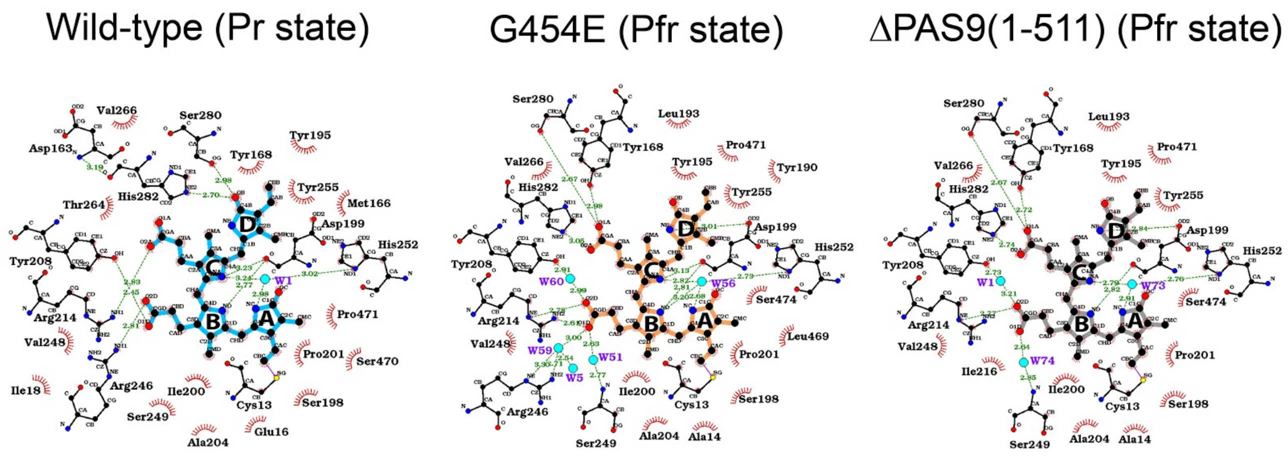


**Figure S6. BV chromophore interactions at the *Xcc*BphP variants in the Pr and Pfr states.**
BV is shown as sticks with carbon atoms in cyan (wild-type), orange (G454E), or gray [ΔPAS9(1-511)], oxygen atoms in red, and nitrogen atoms in blue. The four BV pyrrolic rings are indicated as A, B, C and D. The most important residues are depicted in sticks with carbon atoms in black, oxygen atoms in red, nitrogen atoms in blue, and sulfur atoms in yellow. The different interactions are shown as follows: covalent bonds, purple lines; hydrogen bonds and their lengths, green dashed lines; hydrophobic contacts, red radiating arcs between the atoms. Structural water molecules (including the pyrrole water molecule) are represented as sky-blue spheres. The residues and water molecules participating are labeled. The atoms from the most relevant residues and from the BV are named. The schematic diagrams of the protein-ligand interactions were generated using LIGPLOT v.4.5.3 from the EBI web server (https://www.ebi.ac.uk/thornton-srv/software/LIGPLOT/).

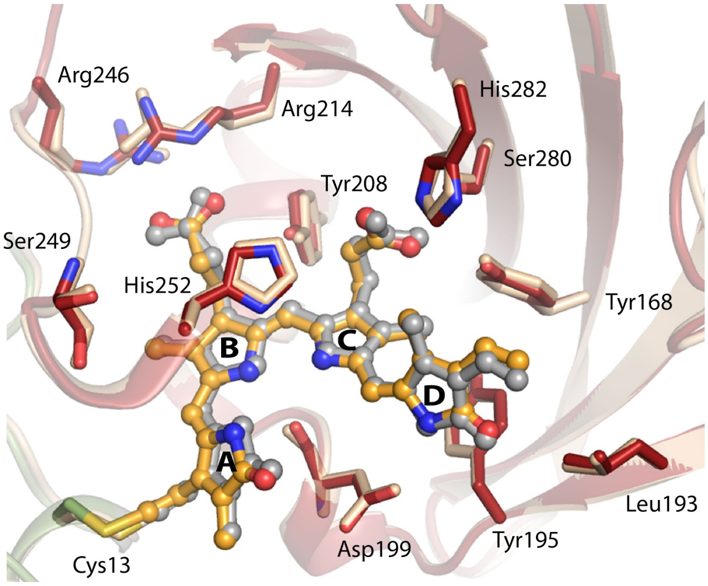


**Figure S7. Chromophore-binding pocket of the Pfr structures.**
Structural contrast between the BV binding pockets from ΔPAS9(1-511) (wheat) and G454E (domains colored according to **Figure 2**). The BV molecule is shown as capped sticks with carbon atoms in grey [ΔPAS9(1-511)] or orange (G454E), oxygen atoms in red, and nitrogen atoms in blue. The four BV pyrrolic rings are indicated as A, B, C and D. The most relevant residues are depicted as sticks; the color code for G454E is carbon atoms in green (within the PAS2 domain) or red (within the GAF domain), oxygen atoms in red, and nitrogen atoms in blue; the ΔPAS9(1-511) residues are wheat-colored. The thioether linkage between the $C3^2$ atom of BV ring A and Cys13 is colored in yellow. Structural alignments were performed on the GAF domain.

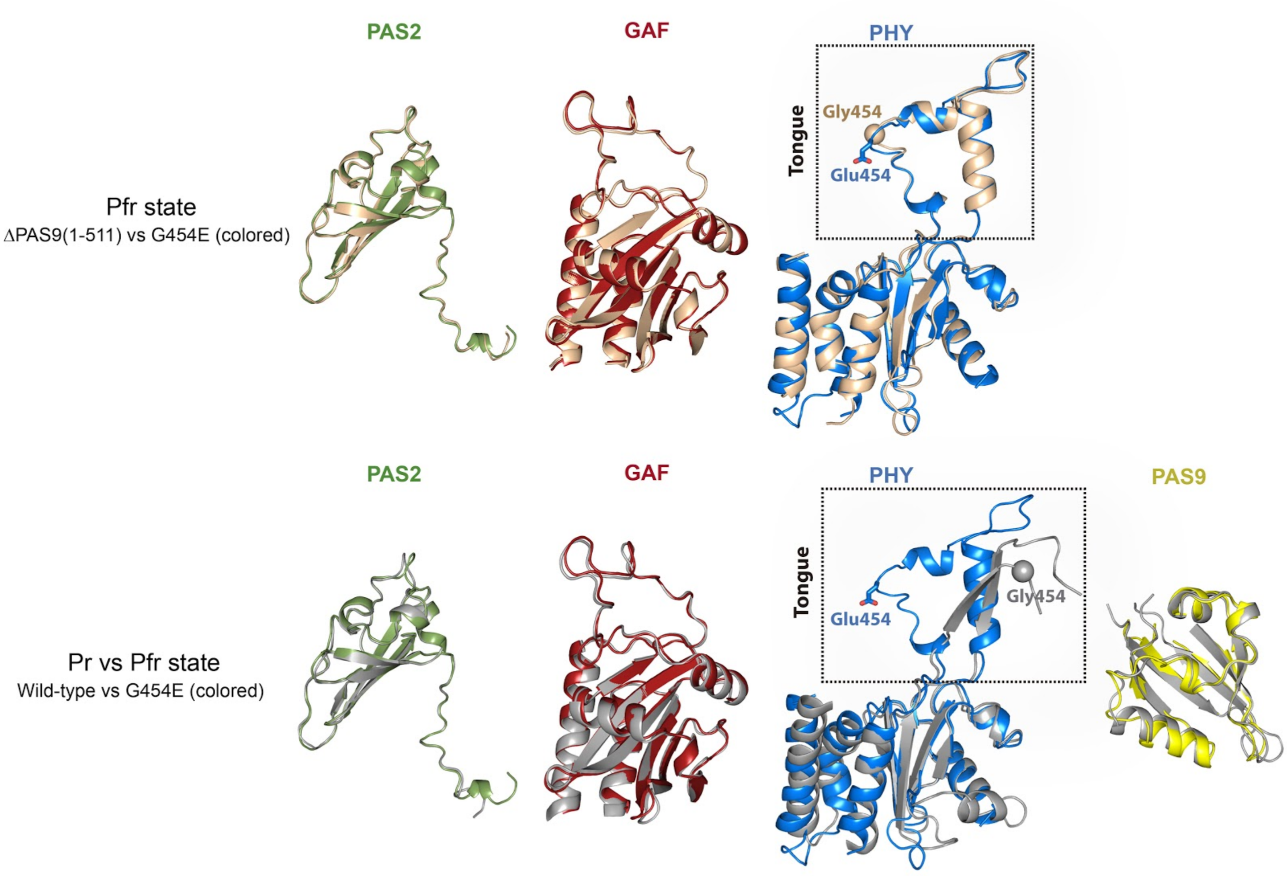


**Figure S8. Structural alignment among the individual domains from *Xcc*BphP variants in the Pr and Pfr states.** *Upper panel:* Self-pairwise alignments between the domains from the Pfr structures of ΔPAS9(1-511) (wheat) and G454E (domains colored according to **Figure 2**). *Bottom panel*: Self-pairwise alignments between the domains from the Pr structure of wild-type (grey), and the Pfr structure of G454E (domains colored according to **Figure 2**). The tongue region in the PHY domain is boxed in a dashed rectangle in both panels. Gly454 and Glu454 are depicted as spheres and sticks, respectively. The different domains are labelled.

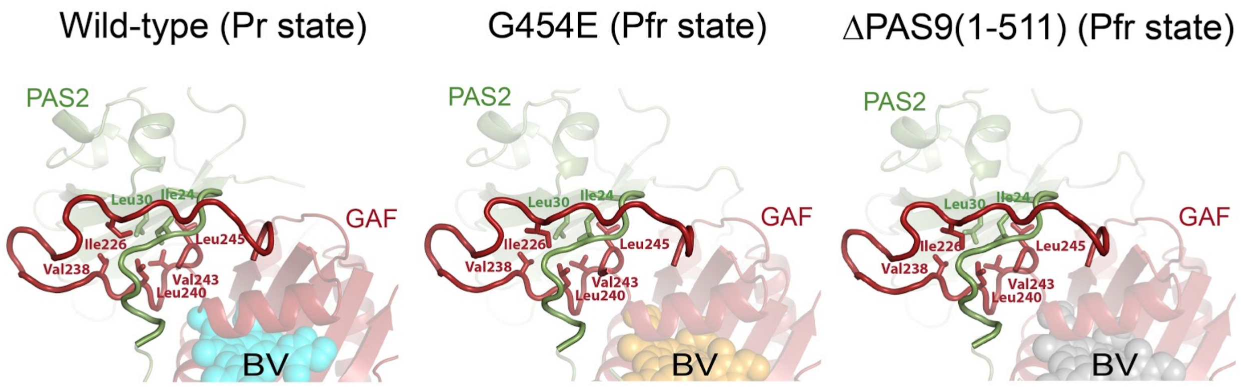


**Figure S9. Figure-of-eight knot from *Xcc*BphP variants in the Pr and Pfr states.**
Ribbon representation of the figure-of-eight knot that crosses over residues between the PAS2 and GAF domains (colored according to **Figure 2**) from the *Xcc*BphP structures in Pr (wild-type) and Pfr (G454E and ΔPAS9(1-511) variants). BV is shown as spheres [cyan, wild-type; orange, G454E; gray, ΔPAS9(1-511)]. The key hydrophobic residues are labeled and shown as sticks.

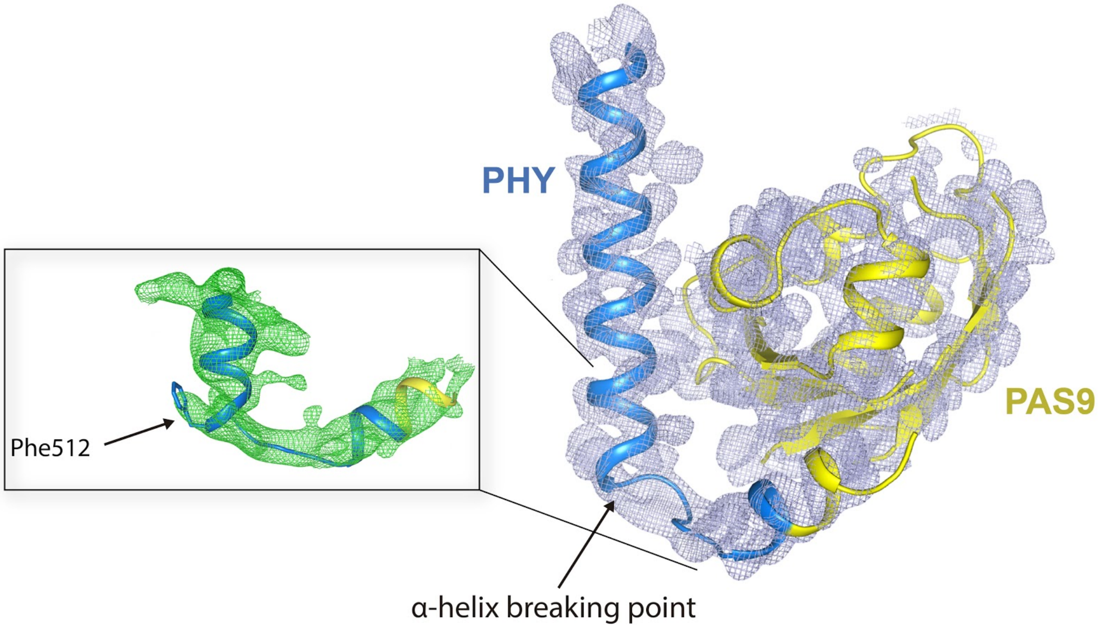


**Figure S10. Electron density maps of the PHY-OM helical linker and the PAS9 domain from the G454E structure in the Pfr state.**

Final $2mF_o–DF_c$ electron density maps (light blue mesh) contoured at the 1.0 σ level around the PHY-OM helical linker and the PAS9 domain (domains colored according to **Figure 2**). The helical "break" is indicated by an arrow. *Inset:* final $mF_o−DF_c$ omit maps (green mesh) contoured at the 2.5 σ level around the break region. Omit maps were generated by removing the residues 504-525 from the model followed by five cycles of refinement in PHENIX (*73*). The location of Phe512 is indicated.

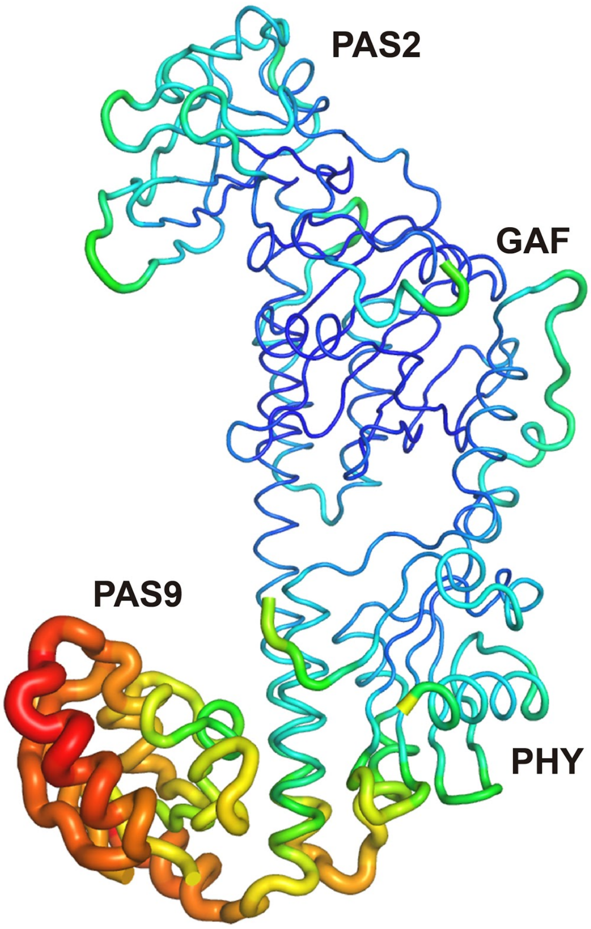


**Figure S11. *B*-factor putty representation of the G454E structure in the Pfr state.**
Putty cartoon representation of the *B*-factor variation on the structure of G454E, colored from low to high (blue to red). The PAS2, GAF, PHY and PAS9 labels are located near their corresponding domains as references.

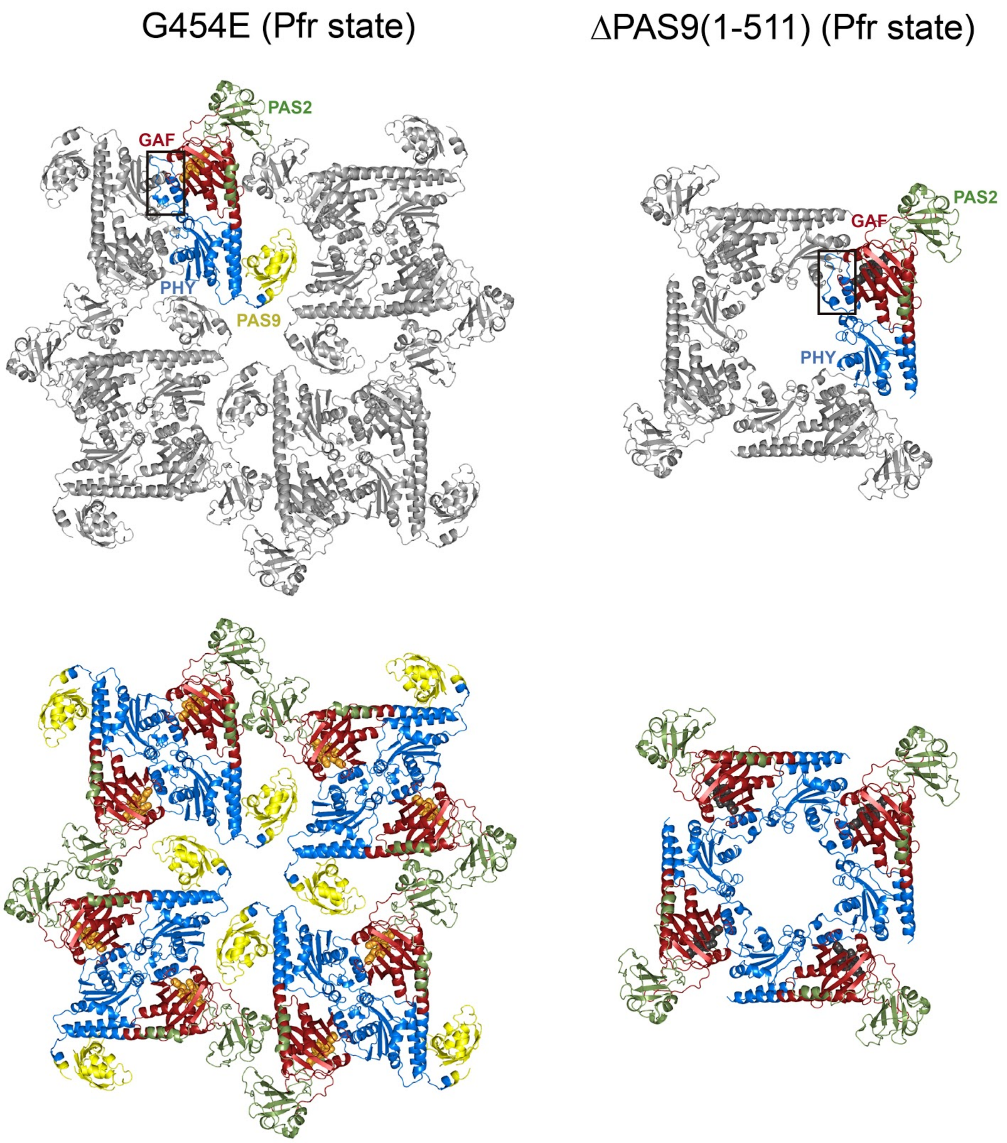


**Figure S12. Crystal packing of the Pfr structures of G454E and ΔPAS9(1-511).**
*Upper panel:* An asymmetric unit is colored according to **Figure 2**, and the symmetry mates are shown in gray. The tongue region is boxed in a rectangle. The different domains are labelled. No crystal contacts are perceived around the tongue region in the crystal packing of ΔPAS9(1-511). *Bottom panel:* All protein units are colored according to **Figure 2.** No crystal contacts are perceived around the PAS9 domains in the G454E crystal packing. In all cases, the crystallographic ***c*** direction is perpendicular to the plane of the image.

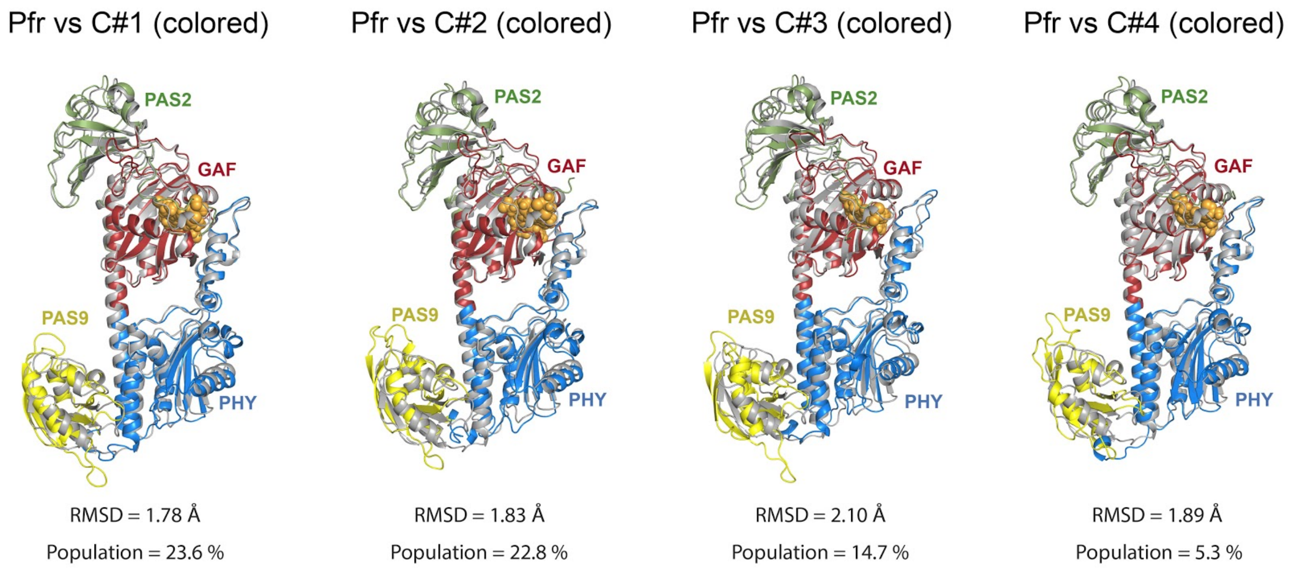


**Figure S13. MD simulations on the monomeric G454E structure in the Pfr state.**
Gallery of the structural alignments between the Pfr crystal structure of G454E (gray) and the MD averaged models (colored according to **Figure 2**, Pfr structure) from the clusters C#1, C#2, C#3 and C#4 obtained from the Pfr structure of G454E. The RMSD (Å) and population fraction (%) of each cluster is indicated. The different domains are labelled.

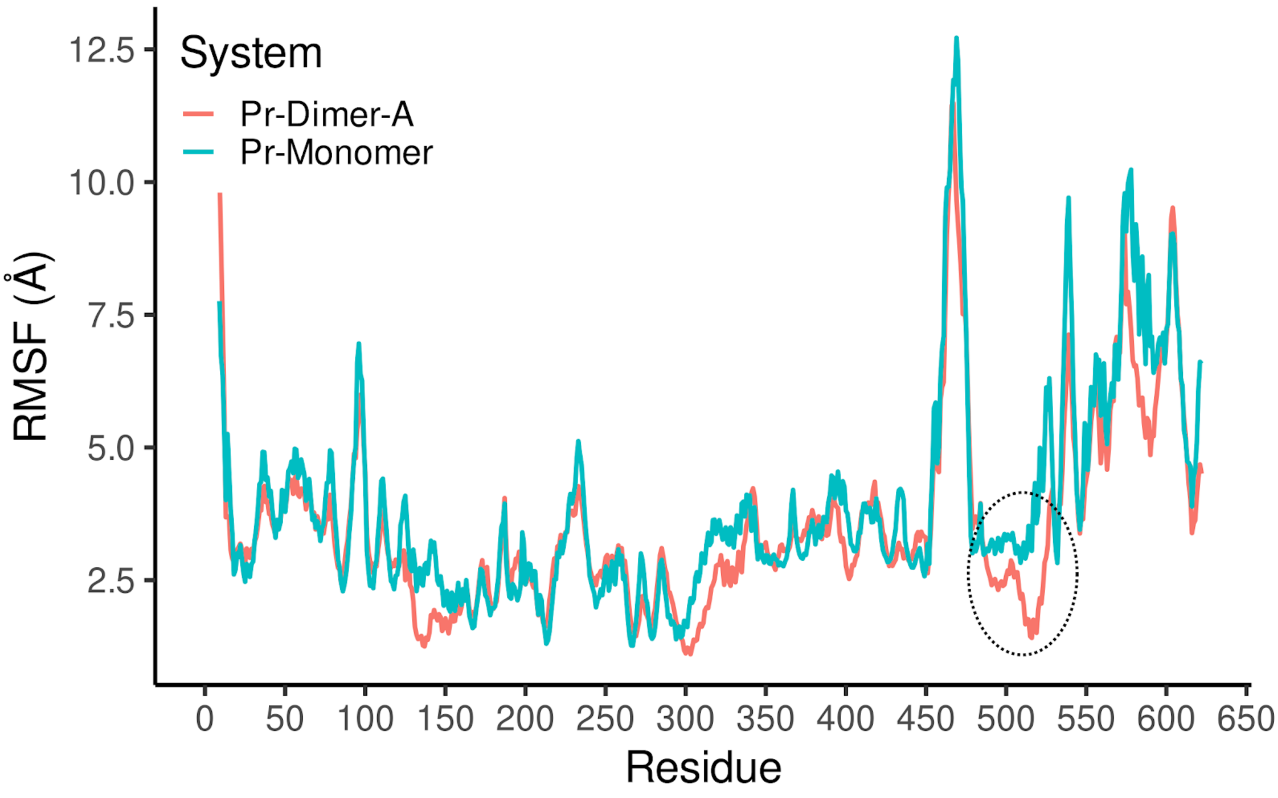


**Figure S14. Average RMSF from MD simulations on the wild-type structure in the Pr state.**
In cyan, the curve corresponding to the Pr monomer while in red, the Pr dimer chain A is shown. A black dashed circle denotes the region around the "break" observed in the PHY-OM helical linker of the G454E Pfr structure.

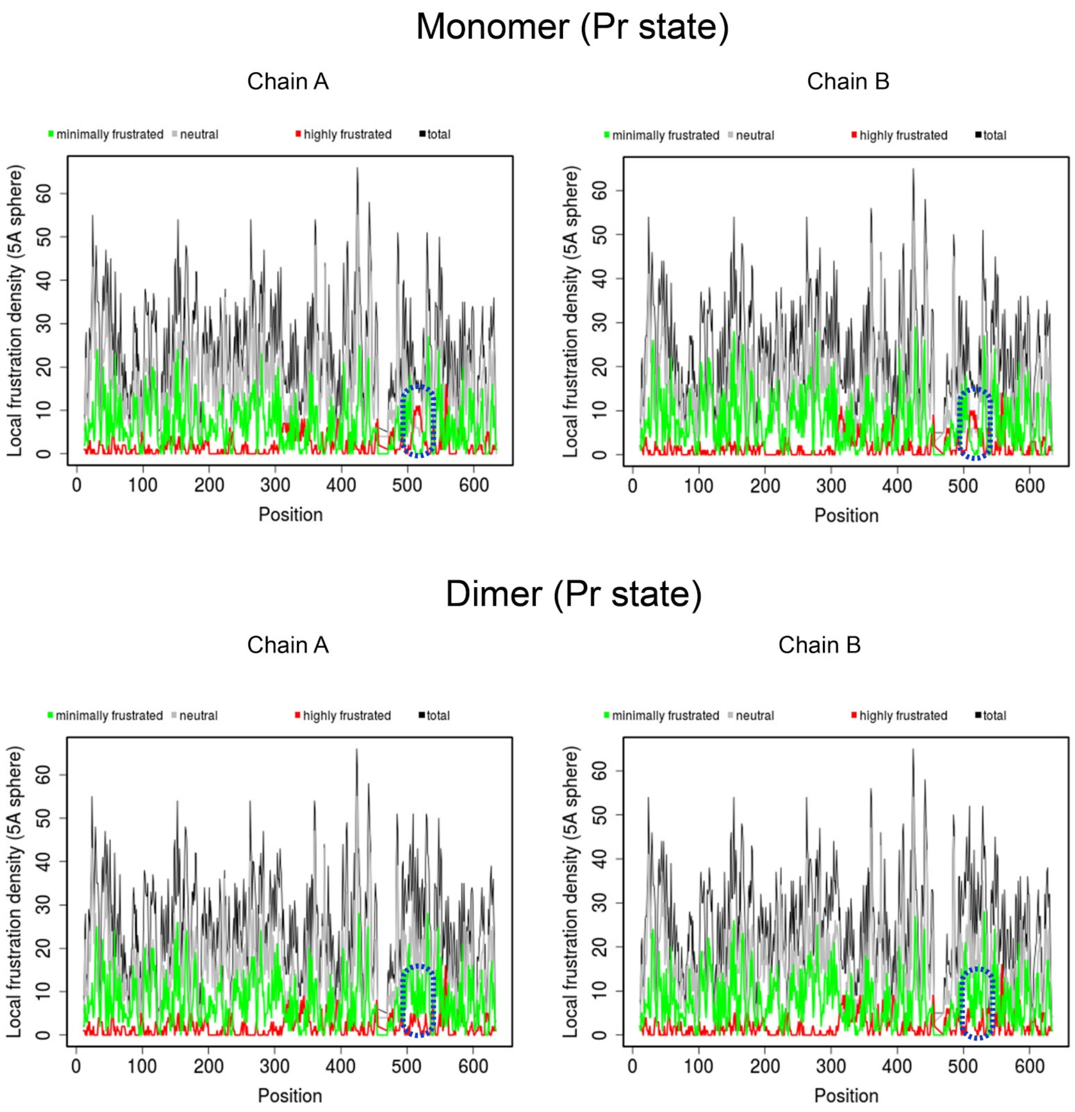


**Figure S15. Frustratograms of the wild-type tertiary and quaternary structures in the Pr state.**
Quantification of the total (black), neutral (gray), minimally frustrated (green) and highly frustrated (red) interactions in the vicinity of each residue (5 Å sphere of each $C_\alpha$), in the Pr monomer chain A and B (upper panel), and in the Pr dimer chain A and B (bottom panel). The PHY-OM helical linker region where the "break" is observed in the Pfr structure of G454E is stressed by a blue dashed oval in both panels. The calculations were performed using the Frustratometer server (*39*) with electrostatics_k = 4.15 (assuming that the environment is totally aqueous).

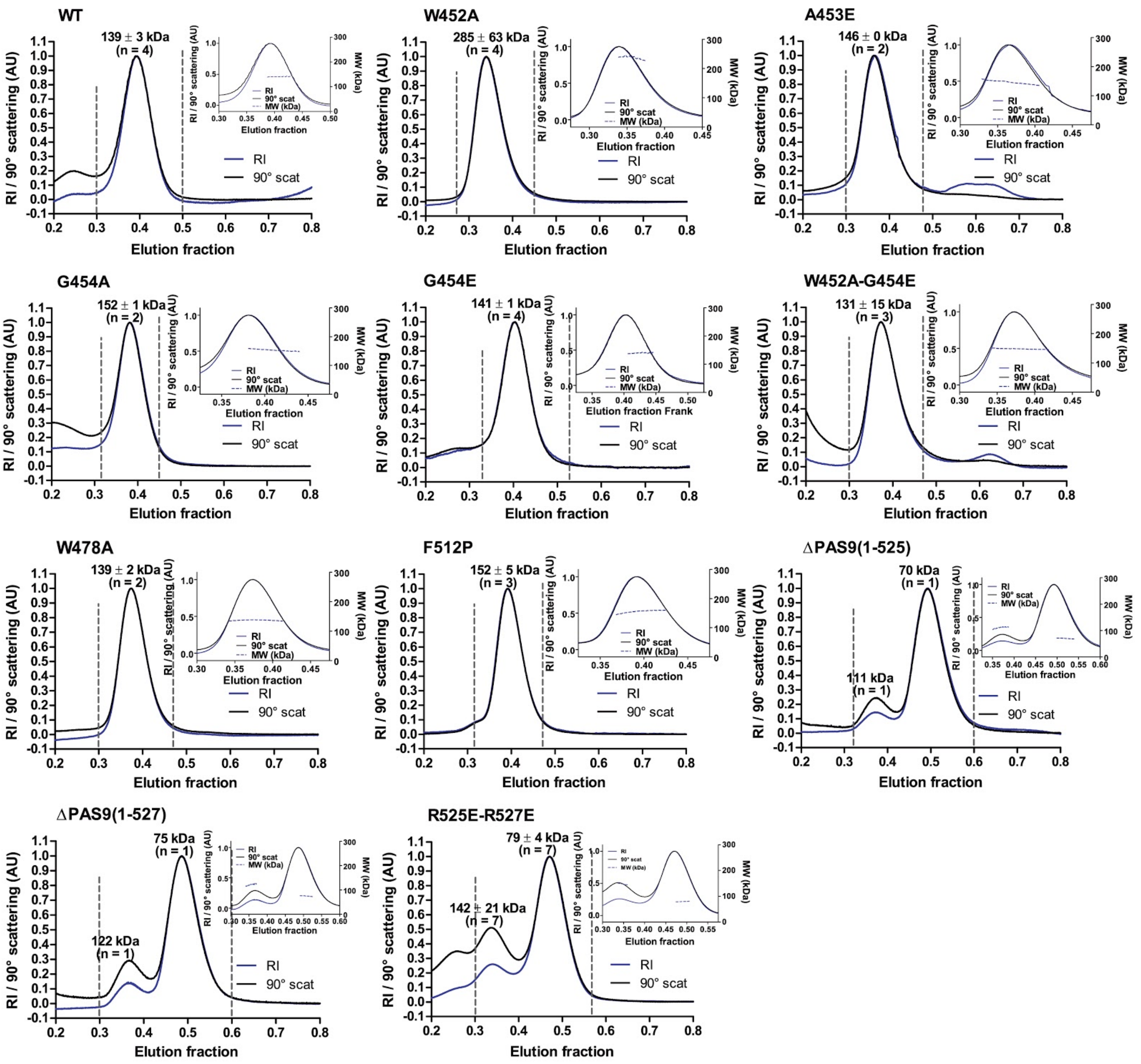


**Figure S16. Light scattering coupled to size-exclusion chromatography of *Xcc*BphP variants.**
The apoproteins were subjected to SEC using a Superdex S200 gel-filtration column. The 90º scattering and the refractive index signals are plotted versus the elution fractions from which the average molecular weights (MW) of the peaks were calculated. The insets show the fraction range corresponding to the peaks and the MW on the right axis. The number of replicates (n) are indicated.

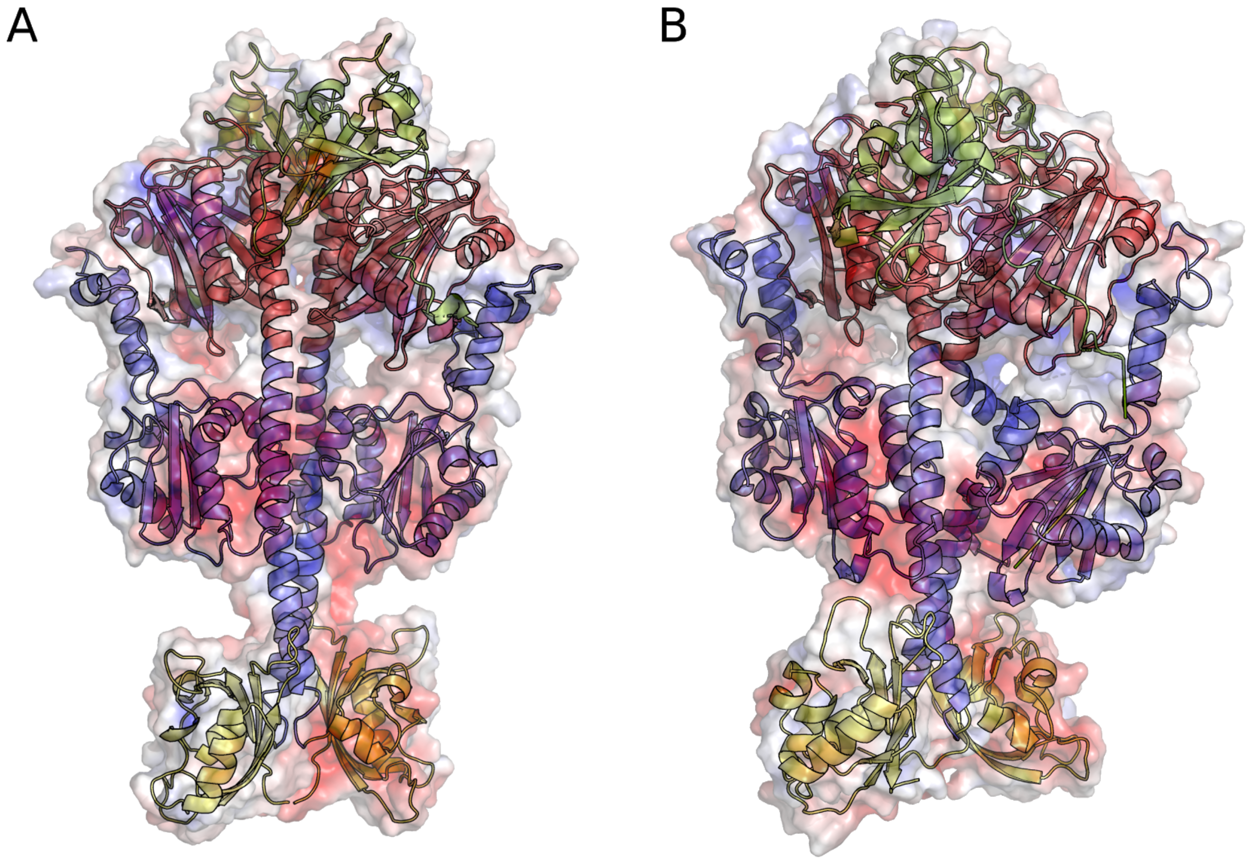


**Figure S17. Projected parallel head-to-head dimer model of the G454E variant in the Pfr state.**
This model was used as comparison to the head-to-tail dimer and to derive values in Table S3. The model was built from the Pr head-to-head dimer conformation and the two PAS9 OMs, in yellow, rearranged to avoid steric clashes as observed in the Pr state. Panel A is the initial structure, and panel B is the final structure after 500 ns GaMD. The dimer interface is conserved with some minor rearrangement of the PAS2 and PAS9 domains. The surface is colored based on the electrostatic values obtained with APBS, ranging from -5, in red, to +5, in blue, units are $K_bT/e_c$.

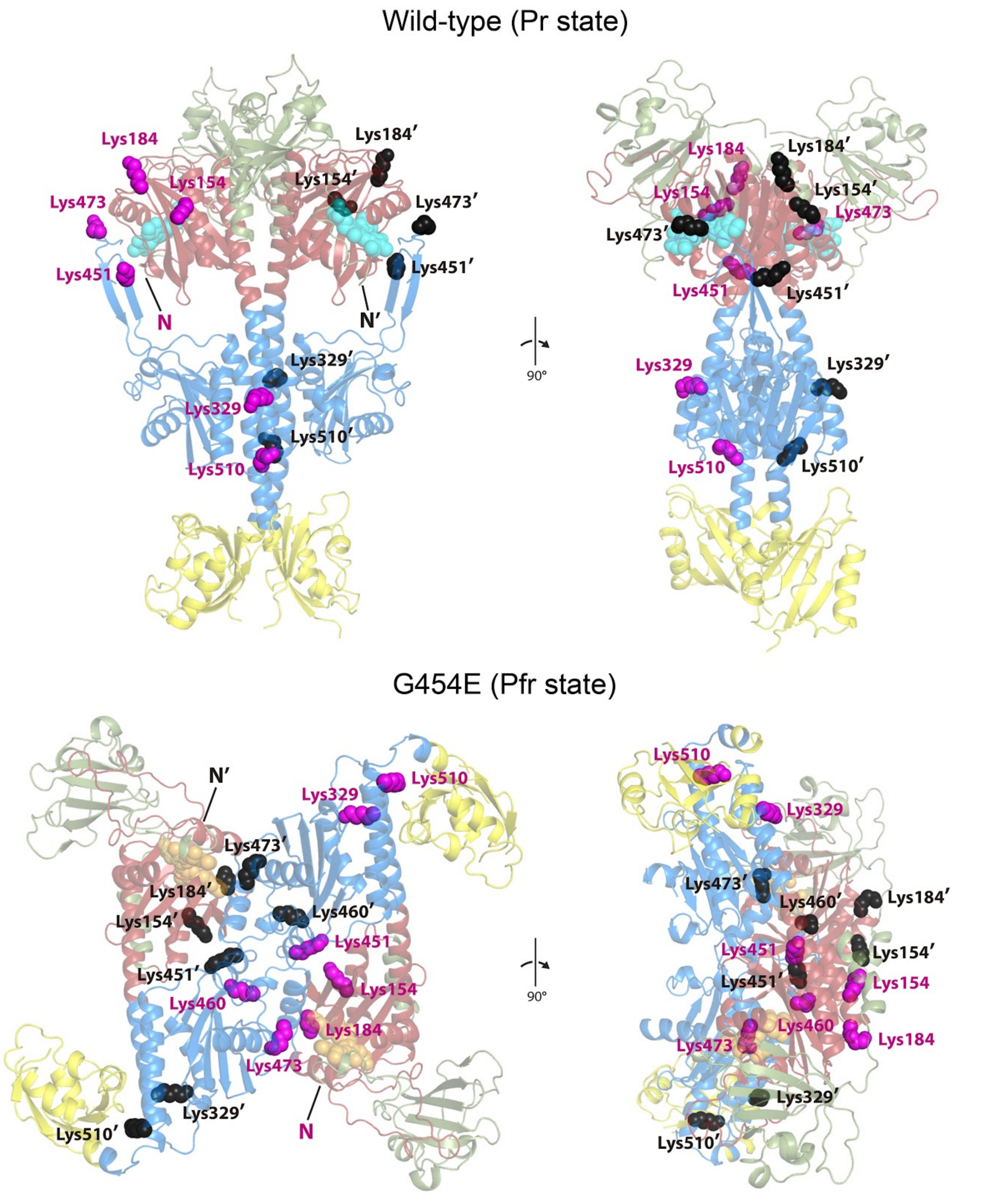


**Figure S18. Lysine residues mapped onto the full-length Pr and Pfr dimer structures.**

The dimeric wild-type *Xcc*BphP Pr and G454E Pfr structures are represented in semitransparent ribbon models with both chains colored according to **Figure 2**. All lysine residues are labeled and represented in spheres color coded by chain (magenta for one chain and black plus prime for the other one). Two pairs of Lys (Lys451/Lys460' and Lys451'/Lys460) are within the cross-linking distance range. The N-termini are indicated and color coded accordingly. Both structures are displayed in two different orientations rotated by 90°.

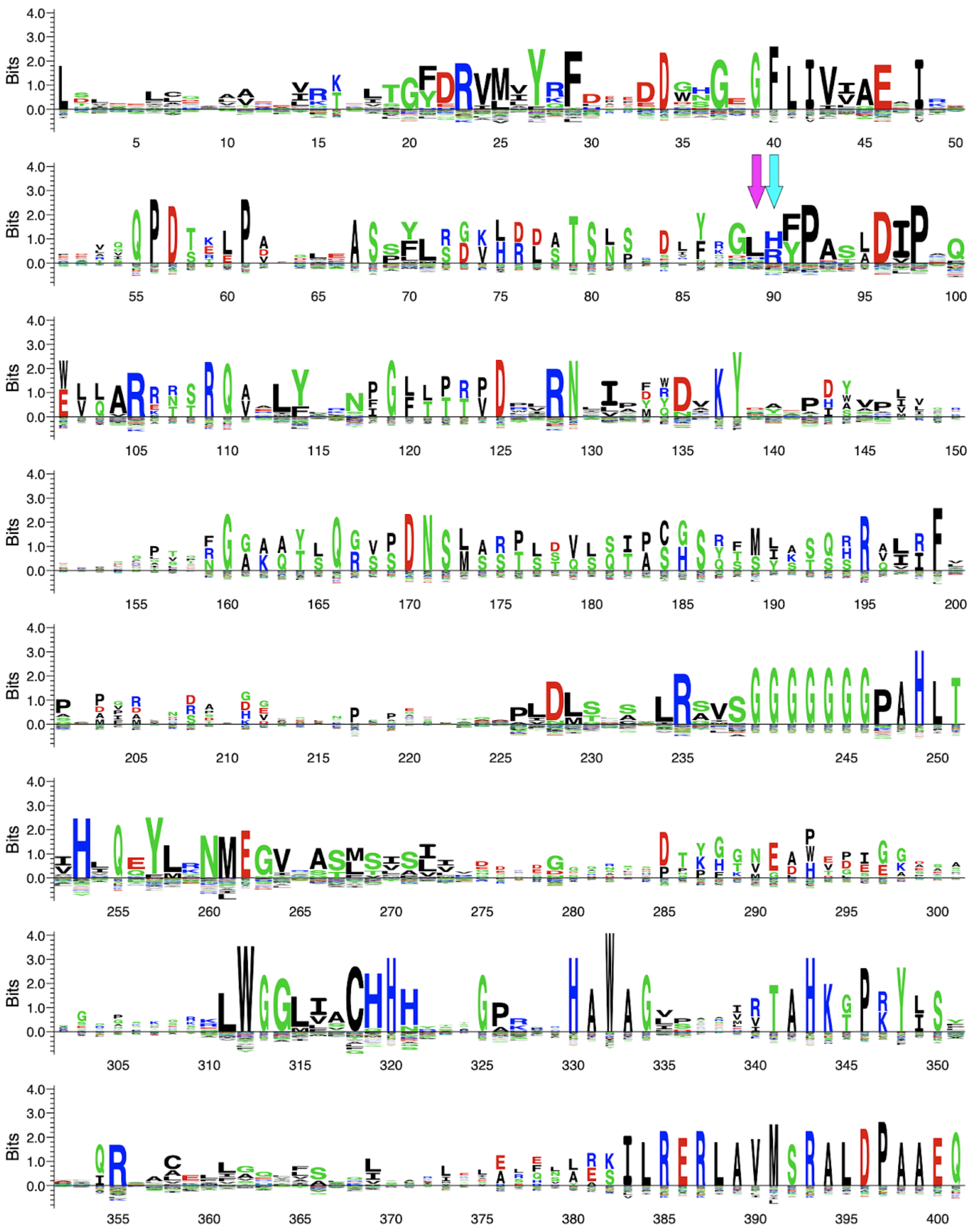

**Figure S19. GAF domain sequence patterns from the phytochrome family.**
The logo was calculated from a GAF domain sequence MSA of 751 non-redundant phytochrome sequences, using the Seq2Logo web server (http://www.cbs.dtu.dk/biotools/Seq2Logo/). The x-axis indicates the residue positions in the MSA. The residue numbering starts from the first residue of the GAF domain MSA. The positions that correspond to the *Xcc*BphP residues Leu193 and His194 are indicated with magenta and cyan arrows, respectively. The y-axis indicates the amount of information for each position in bit units. A score of 4 bits means 100% conservation.

**Table S1. Oligonucleotides used in this study.**

| Name | Sequence (5'-3') | Source |
|---|---|---|
| XccBphP_F | atcatatgcaccatcaccatcaccatagcactgcaaccaacc | This work |
| D177A_F | atgaggagtggaacggcgccatcat | This work |
| D177A_R | tcggcgatgatggcgccgtt | This work |
| H194A_F | aggcctatctcggcctggcgtaccccgcca | This work |
| H194A_R | atgtcgctggcggggtacgccaggccgaga | This work |
| S280V_F | atgcgctgtgggggctgatcgtgtgccat | This work |
| S280V_R | atgcgggctgtagtgatggcacacgatcag | This work |
| W452A_F | atccagcagatcaaagcggccggcaat | This work |
| W452A_R | agctgcggattgccggccgctttgat | This work |
| W452A-G454E_F | atccagcagatcaaagcggccgaaaatccgca | This work |
| W452A-G454E_R | ttggccagctgcggattttcggccgctttgat | This work |
| A453E_F | atccagcagatcaaatgggaaggcaat | This work |
| A453E_R | agctgcggattgccggccgctttgat | This work |
| S574E_F | cccaactcgcgcttgtcgccacgcaaggagttcgatctgtggcagcagacggtgcgc | This work |
| S574E_R | gcgcaccgtctgctgccacagatcgaactccttgcgtggcgacaagcgcgagttggg | This work |
| W478A_F | acgcaagagtttcgatctggcgcagca | This work |
| W478A_R | accgtctgctgcgccagatcgaaactct | This work |
| R501A_F | aatcggcccgcagcctggccgtgctgat | This work |
| R501A_R | atcagttcgatcagcacggccaggct | This work |
| F512A_F | atggagcgcaagcgcgcccagca | This work |
| F512A_R | agcagggtgaagtcctgctgggcgcgctt | This work |
| F512P_F | cgcaagcgcccgcagcaggacttcacc | This work |
| F512P_R | ggtgaagtcctgctgcgggcgcttgcg | This work |
| R525E-R527E_F | ttcaccctgctggaagcctcgctctcagagctggaggatggcgtggccatcatcgagcgcggc | This work |
| R525E-R527E_R | gccgcgctcgatgatggccacgccatcctccagctctgagagcgaggcttccagcagggtgaa | This work |
| ΔPAS9(1-525)_R | taggatcctcagcgtgagagcgaggcttccagc | This work |
| ΔPAS9(1-527)_R | taggatcctcagcgcaggcgtgagagcgaggct | This work |
| pET24a_Gibson_KmR_F | atgagccatattcaacgggaaacg | (*34*) |
| pET24a_Gibson_KmR_R | agcccatttatacccatataaatcagc | (*34*) |

**Table S2. X-ray diffraction data collection and refinement statistics.**

| | WT | G454E | ΔPAS9 |
|---|---|---|---|
| *Data collection* | | | |
| Beamline | PROXIMA-1 | PROXIMA-2A | PROXIMA-2A |
| Wavelength (Å) | 0.9786 | 0.9801 | 0.9801 |
| Temperature (K) | 100 | 100 | 100 |
| Detector | PILATUS 6M | EIGER X 9M | EIGER X 9M |
| Rotation range per image (°) | 0.1 | 0.1 | 0.1 |
| No. of frames | 2400 | 4000 | 1800 |
| Exposure time per image (s) | 0.1 | 0.025 | 0.025 |
| *Indexing and scaling* | | | |
| Cell parameters | | | |
| a, b, c (Å) | 103.53, 103.53, 346.64 | 182.04, 182.04, 40.06 | 164.38, 164.38, 40.44 |
| α, β, γ (°) | 90, 90, 90 | 90, 90, 90 | 90, 90, 90 |
| Space group | P43212 | I41 | I41 |
| Resolution range (Å) | 50.00 – 2.96 | 45.51 – 2.68 | 51.98 – 2.95 |
| Total No. of reflections | 696989 | 290188 | 63381 |
| No. of unique reflections | *40526* | *18944* | *11755* |
| Completeness (%)[a] | 99.6 (98.0) | 100.0 (100.0) | 100.0 (100.0) |
| Redundancy | 17.2 (17.3) | 15.3 (15.5) | 5.4 (5.5) |
| ⟨I/σ(I)⟩ | 15.6 (0.8) | 15.9 (1.9) | 8.8 (2.2) |
| $R_{meas}$ | 0.150 (4.520) | 0.100 (1.479) | 0.140 (0.819) |
| CC1/2 (%) | 100.0 (39.1) | 99.9 (77.1) | 99.4 (75.3) |
| Solvent content (%) | 62 | 47 | 48 |
| Overall B factor from Wilson plot ($Å^2$) | 140 | 107 | 83 |
| No. of chains per a.u. | 2 | 1 | 1 |
| *Refinement* | | | |
| Resolution range (Å) | 49.60 – 2.96 | 45.51 – 2.68 | 51.98 – 2.95 |
| Number of protein atoms | 9524 | 4713 | 3729 |
| Number of ligand atoms | 86 | 43 | 43 |
| Number of water molecules | 2 | 51 | 48 |
| R | 0.206 | 0.196 | 0.185 |
| $R_{free}$ | 0.248 | 0.236 | 0.26 |
| Rms deviations from ideal values (*74*) | | | |
| Bond lengths (Å) | 0.01 | 0.01 | 0.01 |
| Bond angles (º) | 1.2 | 1.2 | 1.2 |
| Average B factor ($Å^2$) | 121 | 105 | 84 |
| MolProbity validation (*62*) | | | |
| Clashscore | 8.09 | 4 | 8.6 |
| MolProbity score | 2.88 | 2.41 | 2.91 |
| *Ramachandran plot* | | | |
| Favored (%) | 90.9 | 93.4 | 91.3 |
| Allowed (%) | 7.3 | 5.1 | 7.2 |
| Disallowed (%) | 1.8 | 1.5 | 1.5 |
| *Protein Data Bank deposition* | | | |
| PDB entries | 6PL0 | 7L59 | 7L5A |

[a] Values for the outer shell are given in parentheses: WT, 3.13-2.96 Å; G454E, 2.81-2.68 Å; ΔPAS9(1-511), 3.13-2.95 Å

**Table S3. Average values of backbone RMSD, number of formed salt bridges and H-bonds, configurational and binding energy in the Pfr quaternary assemblies.**

| | Crystallographic head-to-tail | Projected head-to-head |
|---|---|---|
| RMSD (Å) | 7.095 | 6.005 |
| Salt Bridges | 206 | 215 |
| H-bonds | 331 | 336 |
| Configurational E (kcal · $mol^{-1}$) | -425311 | -427434 |
| Binding E (kcal · $mol^{-1}$) | 21.9 | 79.3 |
